# Point Mutations in SARS-CoV-2 Variants Induce Long-Range Dynamical Perturbations in Neutralizing Antibodies

**DOI:** 10.1101/2021.08.13.456317

**Authors:** Dhiman Ray, Riley Nicolas Quijano, Ioan Andricioaei

**Affiliations:** Department of Chemistry, University of California Irvine, Irvine CA 92697; Department of Physics and Astronomy, University of California Irvine, Irvine CA 92697

## Abstract

Monoclonal antibodies are emerging as a viable treatment for the coronavirus disease 19 (COVID-19). However, newly evolved variants of the severe acute respiratory syndrome coronavirus 2 (SARS-CoV-2) can reduce the efficacy of currently available antibodies and can diminish vaccine-induced immunity. Here, we demonstrate that the microscopic dynamics of neutralizing monoclonal antibodies can be profoundly modified by the mutations present in the spike proteins of the SARS-COV-2 variants currently circulating in the world population. The dynamical perturbations within the antibody structure, which alter the thermodynamics of antigen recognition, are diverse and can depend both on the nature of the antibody and on the spatial location of the spike mutation. The correlation between the motion of the antibody and that of the spike receptor binding domain (RBD) can also be changed, modulating binding affinity. Using protein-graph-connectivity networks, we delineated the mutant-induced modifications in the information-flow along allosteric pathway throughout the antibody. Changes in the collective dynamics were spatially distributed both locally and across long-range distances within the antibody. On the receptor side, we identified an anchor-like structural element that prevents the detachment of the antibodies; individual mutations there can significantly affect the antibody binding propensity. Our study provides insight into how virus neutralization by monoclonal antibodies can be impacted by local mutations in the epitope via a change in dynamics. This realization adds a new layer of sophistication to the efforts for rational design of monoclonal antibodies against new variants of SARS-CoV2, taking the allostery in the antibody into consideration.

## Introduction

The coronavirus disease 19 (COVID-19) caused by the severe acute respiratory syndrome coronavirus 2 (SARS-CoV-2)^1^ has claimed, as of March 2022, more than 6 million lives worldwide, creating a global pandemic and one of the largest public health crises in human history.^2^ To address this urgent problem, scientists across various disciplines are striving to develop drugs, vaccines, and antibodies against the virus. ^3–7^ The spike proteins, present on the surface of the virus, recognize and bind to the human angiotensin converting enzyme 2 (hACE2) receptor in lung cells and initiate infection.^8, 9^ The receptor binding domain (RBD) of the spike is a key target for drug development and antibody recognition. ^10–16^ A large number of monoclonal antibodies (mAb), as well as the natural antibodies (Ab) generated by the immune system, block infection by binding to the RBD and prevent it from attaching to the hACE2 receptor. Moreover, many of the currently available vaccines, such as the ones developed by Pfizer, ^17^ Moderna, ^18^ and AstraZeneca, ^19^ use the spike protein as their epitope and induce immunity by generating antibodies and memory cells that can potentially recognize regions of the spike protein, including the RBD.

The process of antigen-antibody recognition is of fundamental importance for developing immune response against invading pathogens. Since their discovery in 1975, ^20^ monoclonal antibodies (mAb) found a wide range of therapeutic applications, particularly in the treatment of cancer, chronic inflammation and viral infection.^21–24^ Unlike natural antibodies which have varying sequences, monoclonal antibodies, engineered in the laboratory, have a single sequence, and are thereby tailored to treat specific diseases. They are generated by identical B-lymphocyte immune cells cloned from a unique parent white blood cell. The use of engineered antibodies as drugs has become increasingly effective due to their high binding affinity and antigen specificity, both modulated by the complementarity determining regions (CDR) within the variable heavy (V_H_) and variable light (V_L_) chains. ^25^

Monoclonal antibodies have significant advantages over conventional immunization techniques such as convalescent plasma therapy (CPT), because they reduce the risk of blood-borne diseases, shorten the time needed to generate high-affinity and low-risk antibodies, and allow for variable epitope specificity.^26^ A number of monoclonal antibodies have been reported to neutralize the novel coronavirus and are at various stages of clinical trial, with bamlanivimab and REGN-COV2 having received emergency use authorization (EUA) in the United States of America by the Food and Drug Administration (FDA) as of June 2021.^27^ Molecular structures of multiple monoclonal antibodies in complex with the spike protein of SARS-CoV-2 have now been resolved, facilitating a detailed structural -although not as yet a dynamical-understanding of the antigen recognition process. ^28–35^

The major concern for these treatments is the possibility of the appearance of new variants, given mutations in the RBD region, which can evade immune response and reduce or nullify the action of neutralizing antibodies. Multiple such variants of SARS-CoV-2 have emerged throughout the course of the pandemic, threatening the effectiveness of current therapeutics against the coronavirus. One of the earliest well-documented variant, with mutations in the RBD, was the B.1.1.7 strain, a.k.a. the *alpha* variant. It emerged in the United Kingdom in the second half of 2020 and spread across the entire world vigorously because of its 70-80% higher transmissibility compared to the wild type. ^36^ This variant contains a N501Y mutation in the RBD, which has been attributed to its increased immune evasion capability, although no significant increase in reinfection was observed. ^37^ The next major highly infectious variant was the *beta* variant or the B.1.351 strain, originating in South Africa in October 2020. Apart from N501Y, it has two more mutations in the RBD: K417N and E484K; this mutant reduces the effectiveness of conventional antibody therapy up to 10-fold. ^38^ Later, a new strain (B.1.617) with two RBD mutations, L452R and E484Q, emerged in India and became officially known as the *kappa* variant. A lineage of that strain, B.1.617.2, is named as the *delta* variant by the World Health Organization (WHO). With 50-67% higher infectivity than the *alpha* strain, ^39^ the *delta* variant is designated as a *variant of concern* (along with *alpha* and *beta*) because of its potential to make currently available vaccines less effective in preventing serious illness. Evidence of serious decline in vaccine effectiveness against infection (VE-I) has already appeared, along with the proliferation of the *delta* variant.^40, 41^ Single molecule force spectroscopy experiments, in combination with atomistic simulations, confirmed the direct relation between the change of the receptor binding affinity and mutations in the spike protein. ^42^ The same study also demonstrated the reduction of spike protein neutralization by immunoglobulin G based monoclonal antibodies such as B-R41.

To address this urgent crisis, a number of computational studies have been performed to obtain a molecular level understanding of the effect of mutations in the RBD on its binding with the hACE2 receptor and antibodies. ^43–49^ A recent work, specifically explored the change in binding free energy and molecular interactions between spike RBD and a range of neutralizing nanobodies and monoclonal antibodies. ^50^ Taken collectively, these studies revealed that the residues which got mutated in the variants of concern can play an important role in antibody binding. In addition to directly computing binding affinities, some of these studies proposed the possibility of allosteric communication between the RBD and the receptor or the antibody, and showed that dynamical perturbations within the hACE2 receptor can arise due to its binding to the RBD At the same time, simulations have shown that (i) the dynamics of the glycan coat covering the spike affects antibody binding, and (ii) that antibody binding itself affects the motion of the RBD.^51^

From the results of these in-silico studies, it can be argued that the dynamical changes and allosteric effects can play a key role in determining the relative stability of antigen-antibody complexes or the efficacy of monoclonal antibodies. Although allosteric modulation within the spike trimer has received significant attention throughout the pandemic, the viewpoint that mutations in the spike protein can induce allosteric perturbations inside neutralizing antibodies has remained largely unexplored. This is important because such an allostery can facilitate immune evasion. In the present work, we explore the signatures of RBD mutations in the intrinsic dynamics of the monoclonal antibodies when in conjugation with the spike protein. Such effects can only be manifested in atomistic resolution, making them elusive to most common experimental techniques. To understand these allosteric communication mechanisms in greater detail, we performed extensive all-atom molecular dynamics (MD) simulations to gauge the effect of RBD mutations on the stability, dynamics and the unbinding process of monoclonal antibodies targeted to the SARS-CoV-2 spike protein. Apart from the wild type RBD, we studied four mutant strains with the following mutations: B.1.1.7 (*alpha*) (N501Y), B.1.351 (*beta*) (K417N,E484K,N501Y), B.1.617 (*kappa*) (L452R,E484Q) and B.1.617.2 (*delta*) (L452R,T478K). The two antibodies, B38^31^ and BD23, ^32^ were chosen for the current study because their epitopes contain at least two of the four residues involved in the mutations in the studied variants and also because the new mutations in the *beta* variant have been shown to increase the binding affinity for B38 but decrease it for B23. ^43^ This contrasting property makes it particularly interesting to understand how the molecular interaction, and the allosteric cross-talk with the RBD, differ between these two mAbs.

Both of these antibodies were first extracted from the plasma of COVID-19 convalescent patients.^31, 32^ Each is a dimeric protein comprising a heavy and a light chain (labeled as Chain A and B, respectively, in this manuscript) with clearly distinguishable antigen binding domains (Figure 1). The BD23-Fab is smaller (126 and 107 residues in the heavy and light chain respectively) compared to the B38-Fab (222 and 218 residues in the heavy and light chain respectively). The epitope of the B38 contains the RBD residues 417 and 501 while the residues 452, 484 and 501 are included in the epitope of BD23. The structures of the RBD-antibody complexes as well as the location of the mutations are depicted in Figure 1.

**Figure 1:**
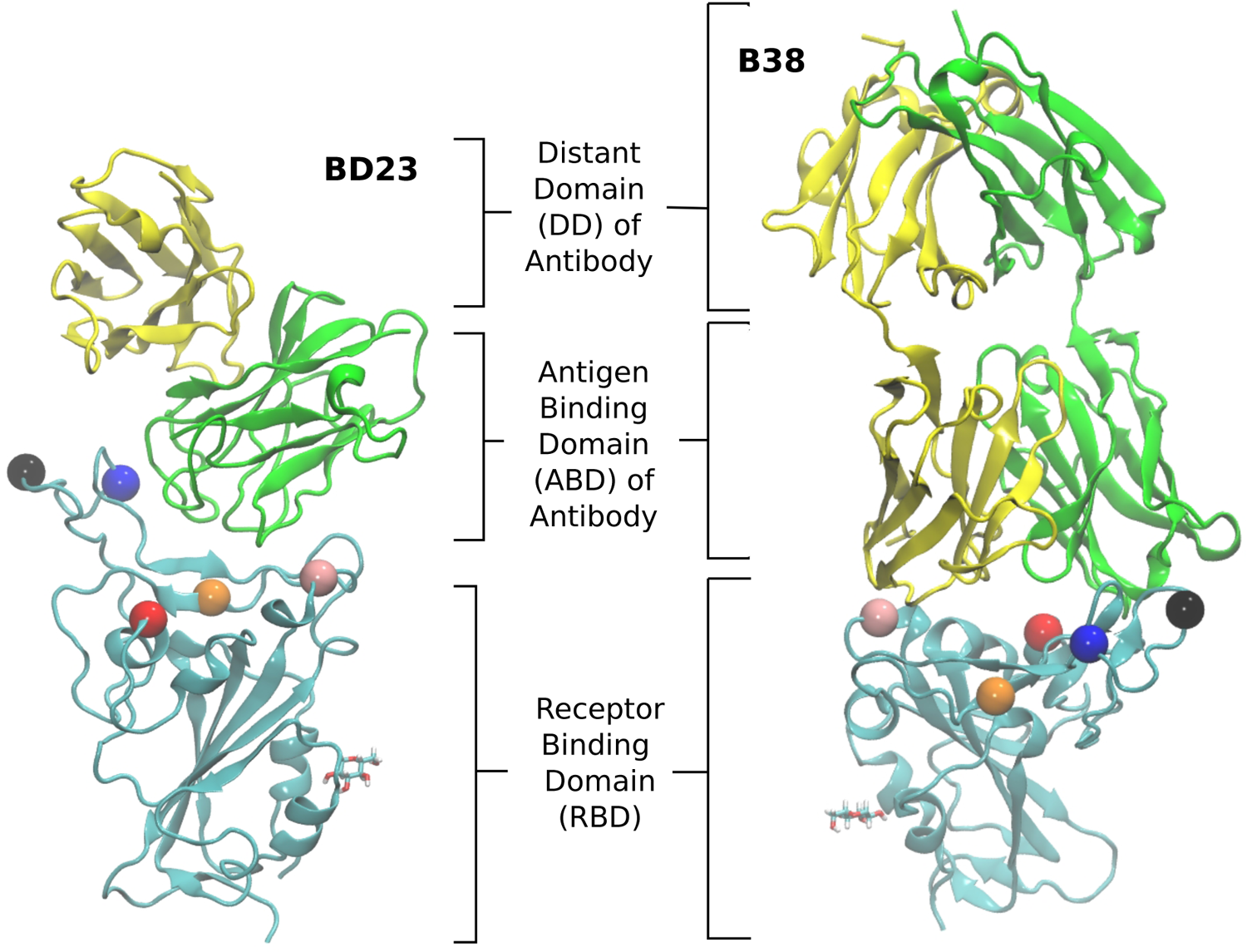
The structure of the RBD in complex with BD23 and B38-Fab, the two neutralizing antibodies studied in this work. Each antibody consists of a heavy (A) and a light chain (B), with chain A in green, chain B in yellow, and the RBD in cyan. The antigen binding domain (ABD) of the Ab is defined as the antibody domain that comes in contact with the RBD; the rest of the antibody is referred to as the distant domain (DD). The glycan moiety on Asn343 is colored in licorice. The residues that are mutated in the studied variants are represented as colored spheres: K417 (red), L452 (orange), T478 (black), E484 (blue) and N501 (pink).

A number of computational studies in the past two years have demonstrated the significance of the glycan shield of the spike protein in receptor binding and immune evasion. ^52–55^ Casalino et al. showed that the roles of glycosilations can go beyond shielding and can alter the conformational dynamics of the spike trimer. ^52^ Such conclusions were reinforced by the work of Sztain et al. where a glycan moiety was identified to work as a gate in allowing the “down” to “up” conformational change in the spike RBD, a mechanistic step, that is essential for SARS-CoV-2 virus to initiate infection. ^53^ As the binding of the hACE2 receptor and neutralizing antibodies generally takes place in the “up” conformation, the effect of glycan moieties in the recognition of the RBD can be less direct. Indeed, Nguyen et al. demonstrated the residue contact signature between the RBD and ACE2 remains unaffected by the presence of the spike glycans. The role of ACE2 glycans, on the contrary, is much more significant in stabilizing te spike receptor complex. ^54^ In the present work, we focused on the role of individual mutations on the dynamics of neutralizing antibodies; so we carefully picked two antibodies whose epitopes inside RBD do not include glycosilated residues. Nevertheless, we explicitly modeled the glycan on Asn343, although it has no interaction with either of the antibodies during the course of the simulation.

We studied the effect of RBD mutations on the dynamics of the antigen-antibody complex from three different angles. First, we demonstrated the relative flexibility of individual residues across the antibody, and identified the structural changes taking place in the complex when the RBD of the WT coronavirus is replaced with each of the mutants. Next, we quantified the change in the mutual information-based cross-correlation between all pairs of residues and constructed a protein graph connectivity network to decipher how are the allosteric information pathways modulated by the mutations in the RBD. Finally, we studied the change in the dissociation mechanism of the antigen-antibody complex for all combinations of the mutant RBDs using non-equilibrium pulling simulations, and rationalized our finding based on arguments involving enthalpy and hydrogen bonding. From the diverse dynamical perturbations observed in these systems, we delineated a few common themes of how the mutations in the SARS-CoV-2 antigen can impart local and global effects through the antibody structure and potentially impair the neutralizing efficacy of monoclonal antibodies. This can occur by changing interaction energies, as well as, via large-scale motions, the conformational entropy of the binding.

## Results

### Modulation of conformational flexibility of RBD-antibody complex

The RBD, with or without the antibody, is a relatively rigid system (RMSD *∼* 1-3 Å) with little conformational variability, likely due to the abundance of beta-sheets in the structure. The antibodies, conversely, are highly flexible with large conformational fluctuations, as indicated by their high root-mean-squared deviation (RMSD) values throughout the production runs (see Supporting Information Figure S3 and S4). Unlike the BD23 system where the entire antibody is highly flexible, in the case of B38, the fluctuations are significantly larger in the distant domain (DD), compared to the antigen binding domains for both chains. The RBD-Ab complexes of both the WT and mutants are also capable of exploring more than one free energy minimum during a period of 0.4 *µ*s, as demonstrated by the projection of their dynamics on the first two principal components (Supporting Information Figure S6 and S7). However, any discernible difference between the dynamics of the WT and mutant complexes is not apparent from the RMSD and PCA analysis.

In contrast, the root-mean-squared-fluctuations (RMSF) of certain residues of both the RBD and Ab do show significant change when comapring the WT to the mutants (Supporting Information Figure S5). To quantify this effect, we computed the ratio of RMSF of each residue for every mutant with that of the WT system. Even in the case of the RBD-only system, the region comprising residues 460-500 shows increased flexibility in the L452R-E484Q variant and decreased flexibility in the N501Y mutant. Interestingly, the E484Q mutation in the *kappa* variant, and the T478K mutation in *delta* variant are present within this region, which also interacts directly with both the B38 and BD23 antibodies. The change in flexibility upon mutation is the highest in the loop formed by residues 470-490, going up or down by a factor of 2-3 in the B.1.617 or the B.1.617.2 strains, respectively. The individually mutated residues show a sharp change in RMSF values, particularly residues 452, 484 and 501 in the respective variants. This effect is also visible in the antibody-bound systems, where the mutated residues show sharp peaks in the RMSF ratio plot (Figs. 2 and 3). Residue 484 shows a contrasting behavior in the presence of B38 and BD23 antibody. When in complex with B38-Fab, K/Q484 and adjacent residues show a large increase in flexibility (*∼*2 times that of E484 in WT), but they become almost twice more rigid when bound to BD23. The RBD of *kappa* variant shows the largest increase in rigidity compared to WT, on the average, when bound to either antibody, but becomes the most flexible of all without antibody. The N501Y, the K417N-E484K-N501Y and the L452R-T478K mutants show increase in overall flexibility when in complex with the antibody. But their RMSF remain virtually unchanged with respect to the WT RBD in the absence of any antibody.

**Figure 2:**
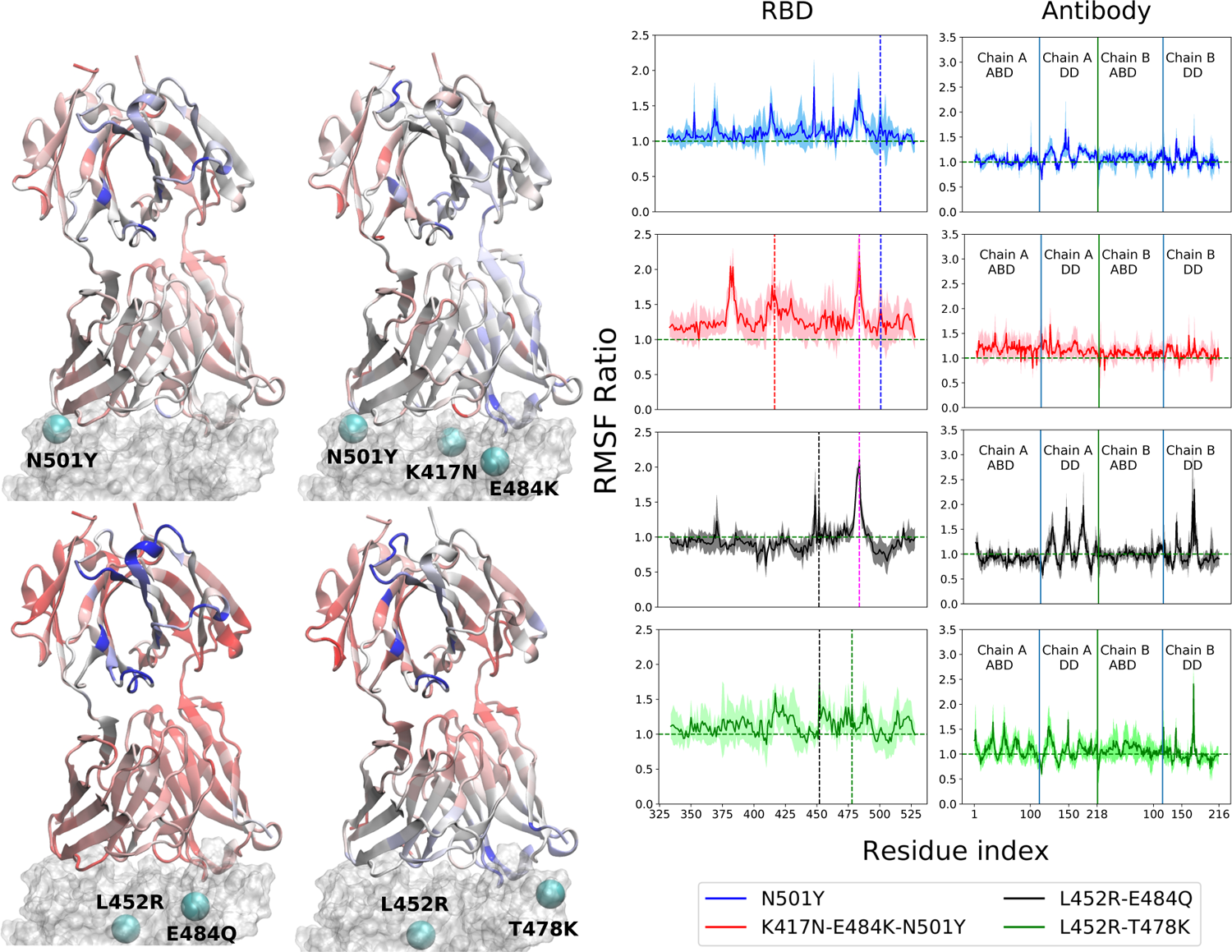
Change of root mean squared fluctuation (RMSF) of the RBD-B38 Fab complex due to the mutations in the variants. The RBD is represented by a transparent surface. The mutated residues are shown as cyan spheres. The antibody residues are colored according to the ratio of their RMSF when bound to a mutant RBD with that of the WT case. Blue color indicates increase in RMSF and red color indicates decrease in RMSF. The residue RMSF ratio is projected on the structure of the B38-Ab complex for *alpha* (N501Y), *beta* (K417N-E484K-N501Y), *kappa* (L452R-E484Q) and the *delta* (L452R-T478K) variants. The RMSF ratio for the RBD and the antibody are shown separately in the right column. See text for the definition of the RMSF Ratio. In the RBD RMSF ratio figures, the location of the mutations are shown as vertical dashed lines.

**Figure 3:**
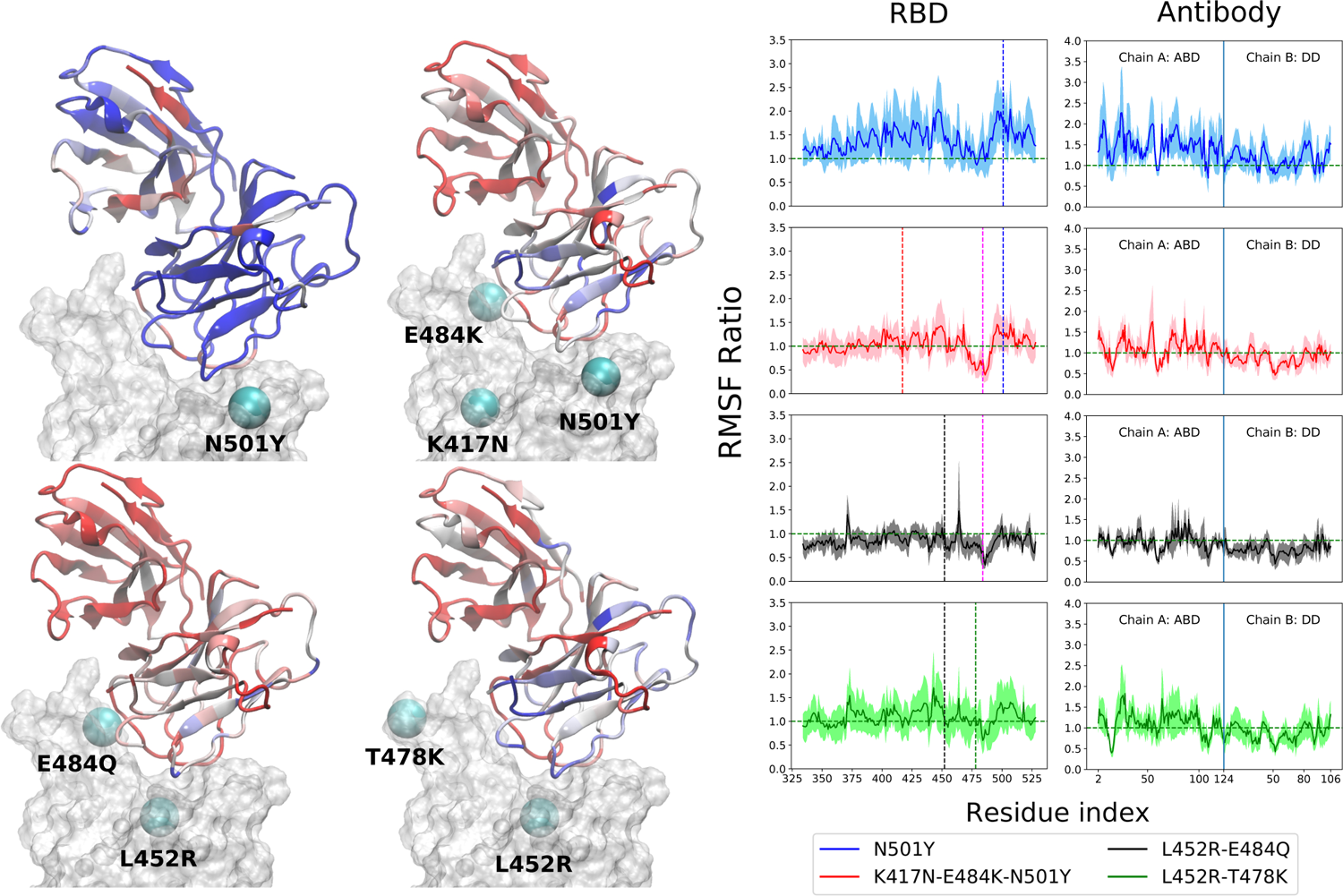
Same as Fig. 2 but for the RBD-BD23 Fab complex.

More interesting is the change in RMSF in the antibody, because this change indicates that mutations in the antigenic epitope can influence the microscopic dynamics of the antibody it binds. The change in the residue-wise RMSF of both the antibodies are, as expected, less dramatic when compared to that in the RBD, likely because these changes arise from a secondary perturbation caused by mutations outside the protein domain (i.e., via non-covalent interactions). Figs. 2 and 3 show the ratio of the RMSF of the antibodies in complex with the mutant RBDs with respect to their WT counterparts. For B38-Fab, the deviation is small (RMSF ratio within 0.5-1.5), except for specific residues in the distant domain, when bound to the L452R-E484Q mutant. However, the changes in flexibility are much more prominent for the BD23 antibody, for which RMSF increases on average in response to a N501Y mutation in the *alpha* variant, but the effect is largely compensated by the K417N and E484K mutations in the *beta* variant, bringing the RMSF down to a level comparable to a WT complex. The L452R and E484Q mutation in the B.1.617 strain causes an increase in overall rigidity of the antibody structure, particularly in the distant domain. The change of antibody RMSF for the *delta* variant case is almost identical to the *kappa* variant, except the antigen binding domain for BD23 shows a slight increase of flexibility in comparison to the *kappa* variant.

### Change in correlated motion and allosteric information flow

Significant changes in the residue-wise LMI-based cross-correlation can be observed throughout the complex (both RBD and antibody) in presence of the RBD mutations (see Supporting information Figure S11 and S13). We particularly focus here on the cross-correlation coefficient between the RBD and the antibody residues, as this can provide a direct quantitative measure of the degree of dynamical coupling between them. In Fig. 4 we depict the relative

**Figure 4:**
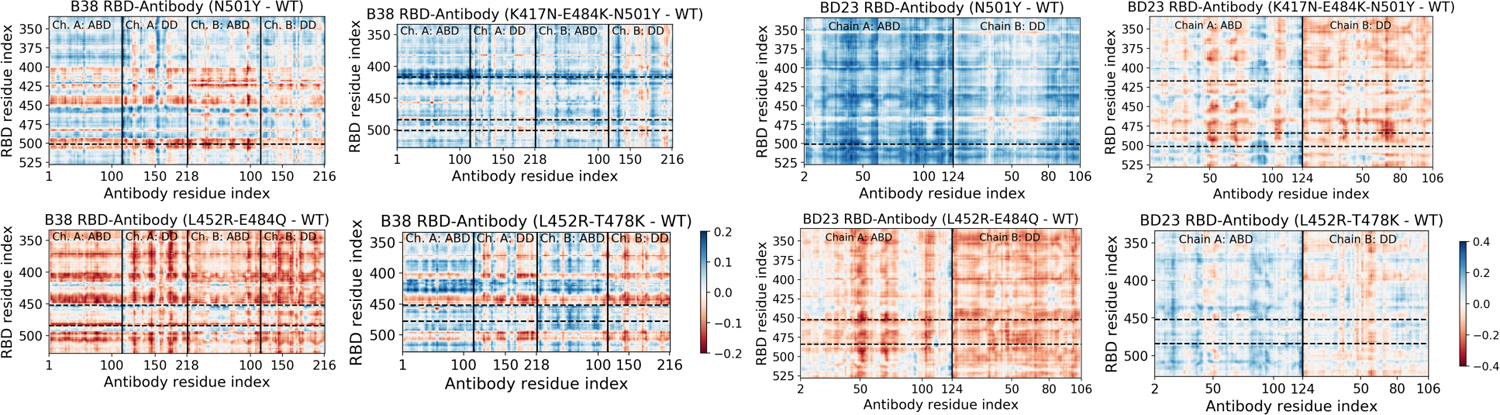
Change in the mutual information based cross-correlation between the antibody and the RBD residues, due to the mutations in the variants. The differences are reported as the cross-correlations of the WT species subtracted from that of the mutant species. The locations of specific mutations are labelled as dashed horizontal lines. The continuous vertical lines separate different domains and chains of the protein (for details, see figure 1).

**Figure 5:**
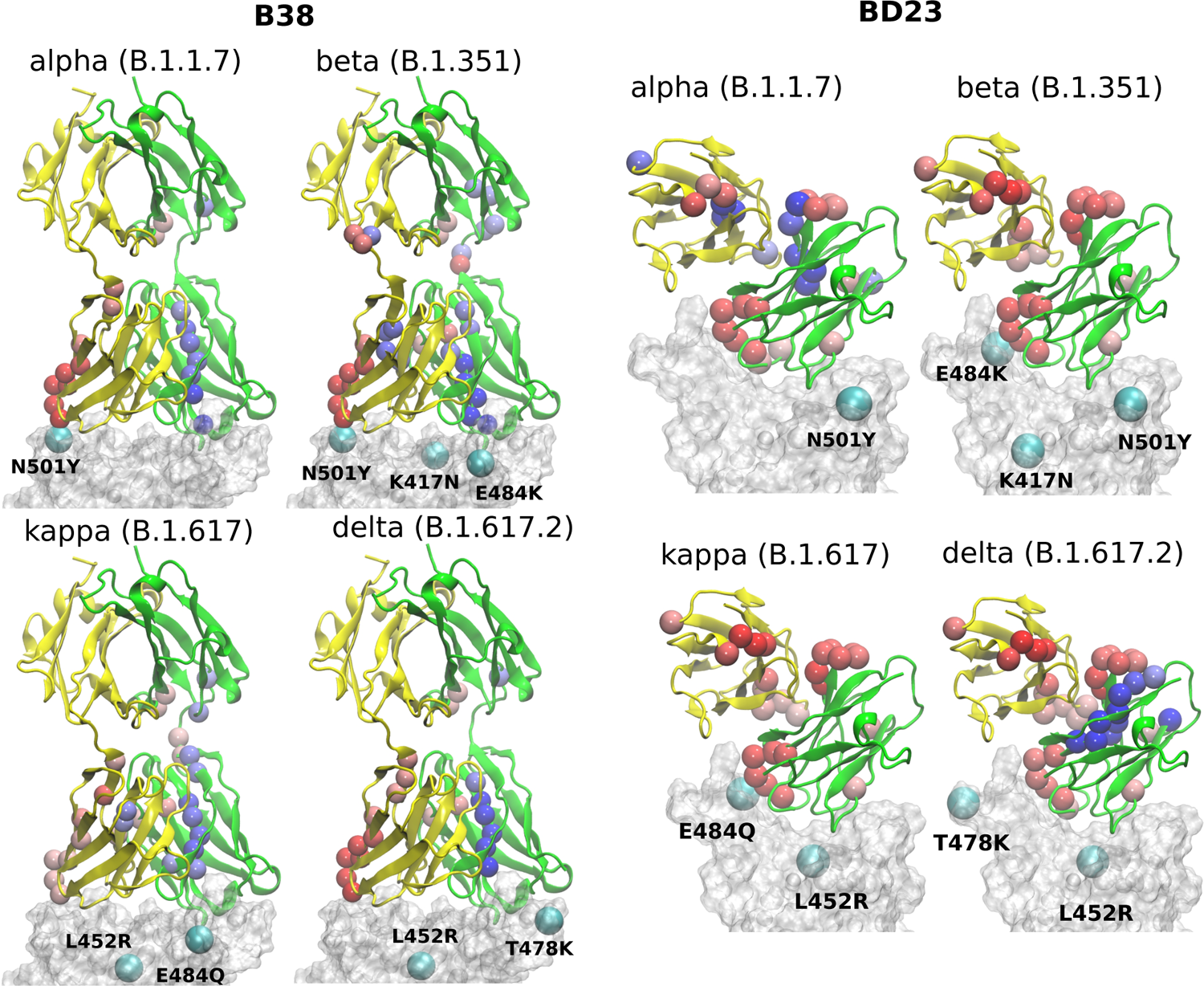
Change of betweenness centrality (BC) of each residue of the RBD-Ab complex due to the mutations in the variants. The RBD is represented by a transparent surface. The mutated residues are shown as cyan spheres. Only residues with a change in BC of greater than 0.005 is shown in colored sphere. Blue color indicates increase in BC and red color indicates decrease in BC.

RDB-antibody cross-correlation for each mutant system, with respect to the WT complex. One of the key features of the B38-Fab is that Ab residues show large change in correlation, particularly with the mutated RBD residues and the residues adjacent to the mutation. For example, in the N501Y variant, many residues of B38 Ab show a decrease in correlation, by 0.1-0.2, with the residue 501 and its nearby RBD residues. More interestingly, this effect is propagated over a long distance, even to the distant domains of the antibody, indicating that single point mutations in RBD can have a strong long range perturbative effect on the dynamics of the antibody bound to the RBD. A similar phenomenon is observed for N417 and its neighboring residues (410-420) in the K417N-E484K-N501Y mutant, the Q484 residue in the case of the L452R-E484Q mutant, and the R452 and its nearby residues (440-455) in the L452R-E484Q and L452R-T478K mutant. Although this effect is less pronounced in the BD23-Fab, the change in cross-correlation coefficients with the mutated RBD residues is indeed observed when, for example, in complex with the K417N-E484K-N501Y mutant. Unlike B38, this effect is less uniform within the antibody, where certain groups of residues (e.g. near residue 50, 70 and 100 in the ABD), show large changes in correlation with the mutated residues as well as for the whole RBD. This can be visualized as vertical patches of red or blue color, as opposed to horizontal patches for B38.

Moreover, there is, on average, an increase in the cross-correlation between the RBD and the B38 antibody, going from the *alpha* variant to the *beta* variant and vice versa in BD23. Previous computational studies have established that B38 binds more weakly with the spike of the *alpha* variant compared to the *beta* variant, but BD23 binds at least twice as strongly to the *alpha* variant than to the *beta* variant.^43, 45^ This observation indicates that the mutual information cross-correlation and the binding affinity between the monoclonal antibodies and the spike epitopes follow a similar trend.

This opposite effect in the correlated motion and the binding affinity suggests that there are allosteric communication paths between the RBD and antibody that follow different routes in the two antibodies. To probe these pathways, we built a protein graph-connectivity network model, treating each residue as the node and the negative logarithm of the cross-correlation between the residues as the weights of the edges of the graph. We computed the betweenness centrality (BC) measure, a metric which computes the relative importance of each node in the information flow pathway. We then computed the residue wise difference of BC in each mutant relative to the WT system. Changes in BC point us to the residues which either gain or lose importance in the allosteric information flow pathway in the protein. In B38-Fab, significant changes in the BC metric are primarily limited to the residues of the antigen binding domain, showcasing a situation where local correlation propagates through the amino acid sequences, leading to long range dynamical changes. Particularly, we see a large increase in the BC value for a few consecutive residues in the region of residues 92-98 in chain A, and a large decrease in the BC for residues 23-30 in chain B, for all the mutant species. This imbalance is primarily caused by the differential interaction of the 501 residue in its WT (N501) compared to its mutated form (Y501). The change in BC in chain B is reduced by a significant amount when the N501Y mutation is removed, for example in the L452R-E484Q variant. This effect can be a result of the reduced interaction energy of Y501 with the B38 mAb as well as the loss of the hydrogen bond between the RDB N501 residue and the S30 residue of chain B of B38, although, there is an overall increase in the hydrogen bond occupancy for Y501 as the lost bond is replaced by a hydrogen bond with S67. In essence, the allosteric information flow through the B38 antibody follows a re-wiring mechanism. The mutations switch off the information flow pathway along the residues 23-30 in chain B and increases the information traffic along residues 92-98 in chain A. This effect is highly asymmetric and location specific, and leads to a drop or rise of the cross-correlation of all antibody residues with particular mutated residues in RBD.

Unlike the B38, in BD23-Fab a large fraction of the RBD-mAb interface (specifically the region containing residues 97-107 of the ABD) loses centrality, along with some isolated, distant domain residues; this theme is common across all mutants. Reduction of BC is also observed for residues 40-50 in the ABD and residues 80-90 in the DD, indicating the possibility of long-distance propagation of an allosteric signal. However, the dynamical effect is different from that in the B38, as the perturbation is not localized in the vicinity of any specific mutation. The same set of residues experiences a decrease in the centrality measure, irrespective of the specific location and the nature of the mutation (except for the *alpha* and *delta* variants, where some residues show a gain in BC, particularly the ABD 45-50 and DD 85-90). This explains qualitatively the observation of the mutation-specific long distance correlation changes in the case of B38 (horizontal correlation changes in Fig. 4). Contrarily, in BD23, specific residues lose or gain correlation with the majority of the RBD residues and not particularly with the mutated ones (see the vertical correlation changes in Fig. 4).

Taken together, these findings provide fundamental insight into the molecular mechanism of how mutations in the spike protein of SARS-CoV-2 variants impact antibody recognition and the stability of the antigen-antibody complex. Not only do these observations bolster the necessity to develop vaccines against mutated or alternative epitopes, but also focus attention on specific regions of the antibodies that can be mutated or modified to engineer a more robust neutralizing antibody, capable to act against potential future variants of the virus.

### Dissociation dynamics and interactions at the RBD-Antibody interface

The observation of allosteric effects within the antibody structure resulting from mutations in the spike also motivates further investigation into the role of mutations in antibody binding and unbinding processes. Considering the difficulty of simulating the diffusion-assisted binding of the antibody in a biologically-accurate environment, we only investigated the possible alteration of the antibody detachment mechanism due to the mutations in spike variants. This provides an alternate angle to understand the role of antigen mutations in the stability of the antigen-antibody complex from a more dynamical perspective. From multiple steered molecular dynamics simulations (SMD), we elucidated the unbinding mechanism of the monoclonal antibodies from the spike RBD. For detailed inspection, we picked the pathway that corresponds to the lowest amount of non-equilibrium work for each RBD variant and mAb combination, as it would contribute the most to the real unbinding process. For all variants and both the antibodies, the loop consisting of residues 470-490 is found to play an important role in the unbinding process. It is the last point of contact between the RBD and the antibody in the lowest work dissociation pathway for every system, and it functions like an anchor to adhere to the antibody before a complete detachment occurs. A recent computational study revealed that this loop region also functions like an anchor while binding to the hACE2 receptor. ^56^ Unsurprisingly, one of the key mutating residue (E484) is situated in this region of the RBD, and, when not mutated, it is often the last RBD residue that forms a hydrogen bond with the dissociating antibody. But in the K417N-E484K-N501Y and L452R-E484Q variants, this residue is mutated and no longer forms the “last” hydrogen bond to anchor the departing antibody. Instead, other residues from the same loop, primarily S477, which has a hydroxyl side-chain, perform this role. We also observed A475 to form the “last” hydrogen bond for the L452R-E484Q RBD with B38 antibody, with its main chain hydrogen bond donor group (See supporting information Figure S24). Although the 484 residue is not mutated in the *delta* variant, the last hydrogen bonds between the *delta* RBD and the departing antibody is formed by the A475, S477 and E486 residues.

We computed the average non-equilibrium work required to detach the antibody for all systems, expressed as *−* ln(exp(*−W_i_*)), where *W_i_* denotes the accumulated work in the *i*th SMD run, and () is the arithmetic mean. The average work is heavily dominated by the dissociation path that requires the lowest amount of work. In the limit of an infinite number of pulling trajectories or in the limit of infinitely slow pulling, the average non-equilibrium work will become equal to the unbinding free energy. As of the convergence of average work, we want to emphasize that we do not aim to calculate an accurate non-equilibrium work, instead we merely generate samples of work along a handful of paths. So the average work reported in this work is not meant to be used as an estimate of the free energy (as done, for example, when using the Jarzynski identity^57^ or other fluctuation theorems^58^). To establish the validity of our implicit solvent SMD approach, we performed additional steered molecular dynamics simulations in an explicit solvent environment. While it is true that explicit solvent gives more accurate thermodynamics, when pulling a ligand through water the rate of pulling must be accounted for. ^59^ Implicit solvents permit instantaneous (“adiabatic”) rearrangement of the “water”, while explicit water imposes a limit to the speed with which the ligand can be pulled (i.e., slower than the structural relaxation of the explicit water molecules). As a consequence, the absolute work will be different due to interaction with the water molecules during the pulling. However, the relative differences are similar, as we show in the additional calculations (see Supporting Information). Also, the calculation of uncertainty in exponential averaging of non-equilibrium work can be nontrivial due to the presence of finite statistical bias. ^60^ Nevertheless, we showed a very primitive mean-square-error based uncertainty analysis in the Supporting Information (Fig. S22 and S23). Although we can observe different relative uncertainty across different variants, the absolute values of the variance are not particularly meaningful due to the small sample size. Provided E484 is the key anchoring residue, we expect the non-equilibrium work will depend on how strongly this particular residue or its mutants interact with the antibody.

From the non-bonded energy calculation on the equilibrium trajectory it is apparent that the three mutated residues: E484, K484 and Q484, have little difference in their interaction energy with the B38-Fab antibody, primarily because this residue is not in direct contact with the antibody interface (Fig. 6b). Consequently, the four mutants do not show any discernible difference in the work required for the dissociation of the RBD-mAb complex (Fig. 6c). But, the *alpha* and *beta* mutants require the least amount of work for antibody detachment in comparison to the WT, an effect that can partially be attributed to the N501Y mutation. From non-bonded energy calculation, we see that the N501Y mutation destabilizes the interaction between RBD and B38 by lowering (less negative) the interaction energy (see Supporting information Figure S16). Non-bonded interaction energies have previously been used to understand the impact of mutations in the receptor binding affinity of the SARS-CoV-2 RBD.^42^ Similar to the binding of hACE2 receptor, we observed that the change in interaction energy can also have a significant effect on the binding of antibodies to the RBD For the BD23-Fab, an almost opposite effect is observed. The residue E484, being situated in the RBD-mAb interface, plays a key stabilizing role for the complex via hydrogen bond to the S103 and S104 residues of the Ab. Once mutated, the hydrogen bond propensity of residue 484 diminishes, leading to almost 80 kcal/mol decrease (less negative) in the non-bonded interaction energy with the mAb, in agreement with a recent computational study. ^45^

**Figure 6:**
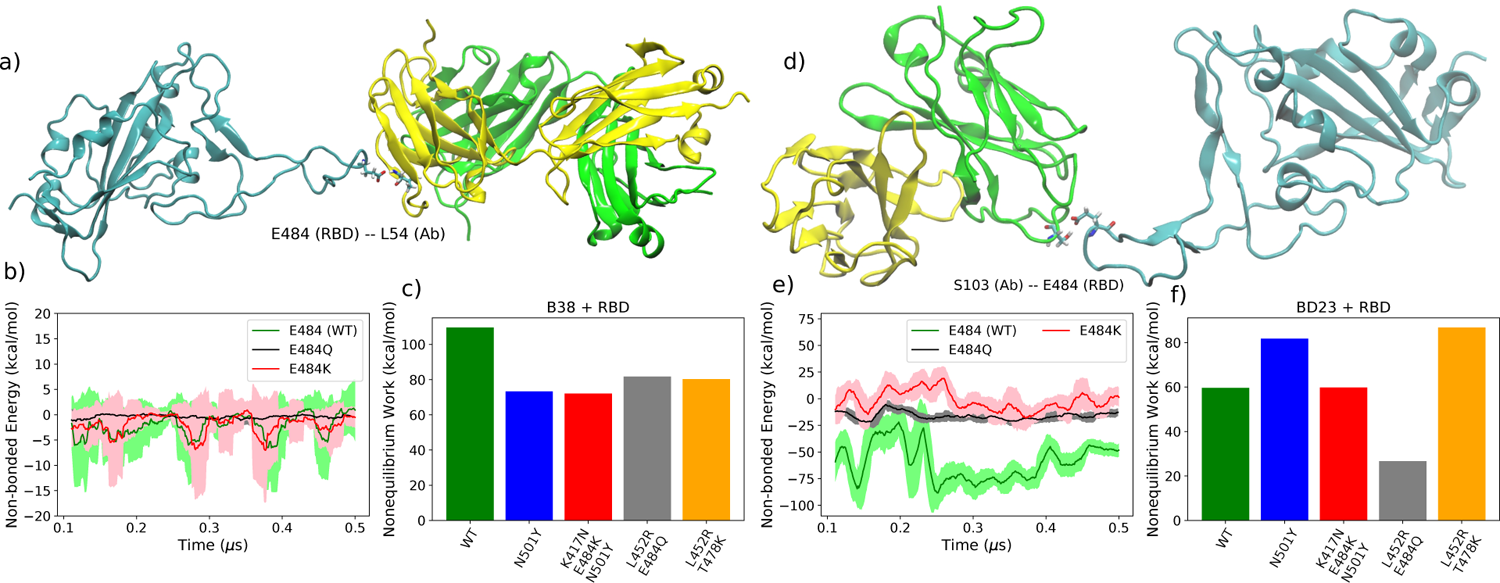
(a) A representative intermediate structure for the B38 antibody unbinding from the RBD from the SMD simulation. Intermediates for all variants are shown in supporting information Figure S24 ans S25. (b) The non-bonded interaction energy between the RBD residue 484 and the B38 antibody from the unbiased simulation. (c) The average non-equilibrium work performed to dissociate the B38 antibody from all variants of the RBD using SMD simulation. Figure (d-f) are identical as (a-c) but for BD23 antibody.

The E484 forms hydrogen bonds with the S104 (26.37% with backbone and 54.19% with side-chain) and S103 (3.85% with backbone and 8.62% with side-chain) residues of BD23 with high occupancy in the WT and with G102 (31.9%), S103 (4.15%), Y32(10.7%), Y110 (3.95%) and Y111 (2.73%) residues of the antibody in the N501Y mutant. In the K417N-E484K-N501Y variant, K484 forms hydrogen bonds only with N92 (6.86%) and Y110 (1.18%) with relatively low occupancy. This effect is clearly reflected in the non-equilibrium work profile, where, going from N501Y mutant to K417N-E484K-N501Y and L452R-E484Q variants, we see a significant drop in the work required for the detachment of the antibody. Conversely, for the *delta* (L452R-T478K) variant, where the 484 residue is not mutated, the E484 forms hydrogen bonds with *>*50% occupancy with the S103 and Y109 residue of the BD23 antibody. This increases the amount of non-equilibrium work required for the dissociation of the RBD antibody complex to a strikingly high level. This is reflected also in the high binding free energy of the BD23 antibody with the *delta* variant RBD.

In contrast, the N501Y mutant, binds more strongly with the BD23-Fab (Figure 6f) compared to the WT, because the non-bonding interaction energy of Y501 is higher (more negative) than that of the N501 residue (Supporting info Figure S17). This effect can be attributed to the increased probability of hydrogen bonding between the RBD and the antibody by the Y501 residue compared to the N501. In the WT system, N501 participates in one hydrogen bond with the D73 residue of the BD23-Fab with negligible occupancy (0.62%). But in the N501Y mutant, the Y501 participates in two hydrogen bonds with the L72 and the Y80 residues of the antibody with significantly higher occupancy (5.77% and 6.27% respectively). The rupture forces of the spike antibody complexes follow a similar trend as the average non-equilibrium work for the WT and the mutants (see Supporting Information Figure S20 and S21).

The relative order of the binding free energies for each RBD antibody pair is also consistent with the average non-equilibrium work necessary for the dissociation of the respective complexes (Fig 7). In agreement with the earlier work, ^43^ the binding affinity of the BD23 with the *beta* variant was observed to be weaker than the *alpha* strain, and vice versa for the B38 Fab monoclonal antibody. This indicates that the arguments presented above, concerning the non-bonded interaction energy and hydrogen bonding propensity, also hold true for the binding free energy of the antigen-antibody complexes. The practical implication of these findings is that antibodies, such as B23, that uses the RBD 470-490 loop as an epitope, are more likely to have significantly different affinity against mutated versions of the virus. Alternatively, fairly consistent neutralization of most variants is possible if antibodies or vaccines are generated using an alternative region of the receptor binding domain as the epitope.

**Figure 7:**
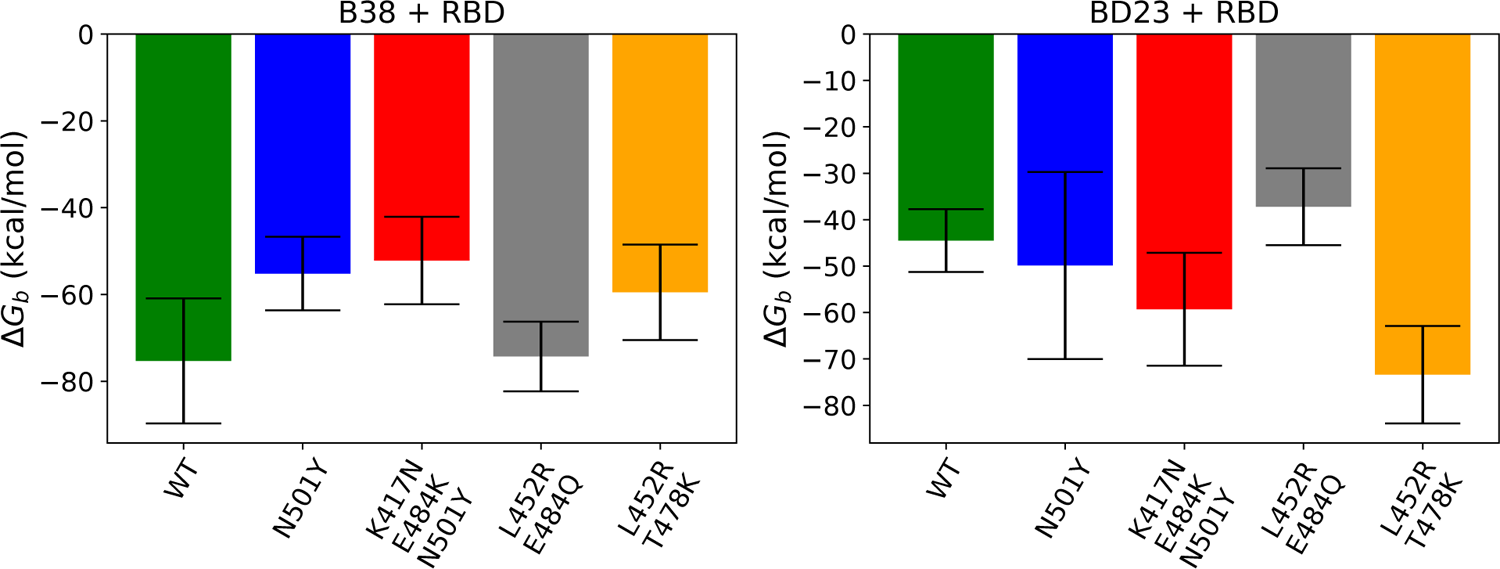
Binding free energy of the spike protein variants with the B38 and BD23 antibody.

## Discussions and Conclusions

The simulations of the interaction between the receptor binding domain of the SARS-CoV-2 spike protein and neutralizing monoclonal antibodies reveals the functional role of point mutations in the new variants in the process of immune evasion. Point mutations perturb the microscopic dynamics of the antibodies and destabilize the antigen-antibody complexes. Key findings of our work are the following. First, the structures of the antibodies are largely flexible and can undergo significant conformational transitions over a timescale of several hundreds of nanoseconds. However, the flexibility of certain residues in the antigen-antibody complex are modified disproportionately when the WT RBD is replaced with one from the mutant strains. The mutated RBD residues (and the residues in proximity with the mutations) show the largest change in flexibility. Particularly, in the absence of an antibody, the loop involving residues 470-490 becomes flexible when a Glu to Lys (or Gln) mutation at the 484 position makes it incapable of forming a stabilizing hydrogen-bonding interaction. While local motions in the antibody occurred next to the mutation site in the spike, there were also significant large-scale collective motions throughout the antibody structure. We speculate that these motions can modulate the free energy of binding via contributions to the conformational entropy. Changes in binding can also result from the overall change of stability of the antigen-antibody complex, arising from the noticeable changes in rigidity observed not only in the antigen binding domain, but also in the distant regions. The effect is more prominent for the smaller antibody, BD23 Fab, where the N501Y mutation drastically increases the dynamic motion across both the domains in the antibody structure. The K417N and E484K mutations reverse this effect to a large extent, and, in the B.1.617 variant, both antibodies become structurally more rigid compared to the WT complex. Because the dynamical perturbations in the *antibody* result from mutations in the *antigen*, we are presented with a striking example of inter-protein allosteric information propagation, which has the potential to affect the evolutionary driving forces on the virus to adapt against neutralizing interventions.

The linear mutual information-based cross-correlation between the antigen-antibody residue pairs is a useful metric that can capture quantitative information about the allosteric information flow. Cross-correlation values, for both the RBD and the antibody, can change significantly due to mutations in RBD, but there is a stark contrast in the manifestation of this change in correlated dynamics in the two antibodies. Unlike B38-Fab, where correlations with specific mutated RBD residues are modified across the entire antibody, such effects in BD23 Fab are fairly localized albeit universal. Certain groups of residues in the BD23-Fab gain or lose correlation with the entire RBD and not just with the mutated residues. The antibody residues impacted by the mutation are conserved for all mutants, but whether they show a positive or negative change in cross-correlation can depend on the nature of the mutation. We previously observed such an effect within the spike protein, ^61^ but now we see that the allosteric effect can be propagated into the antibodies that interact with the spike only via non-covalent interactions. One key observation is that the gain or loss in cross-correlation follows the trend of binding affinity fairly well. This phenomenon behooves one to investigate further, for a wide range of protein-protein complexes, the relationship that inter-residue cross-correlations has with, on one hand, binding free energies. On the other hand, simulation-based correlation paths can put in relationship with experimental measures of allosteric signals transduced within and between proteins. For example, using cryo-EM, Frank and collaborators have delineated a way to retrieve functional pathways of biomolecules from single-particle snapshots. ^62^ Among the various experimental approaches that can probe allostery at the atomic resolution detail is liquid-state NMR spectroscopy. Using it, one can obtain site-specific dynamical information that can be used to probe allostery experimentally, see for example. ^63^ The dynamical parameters measured by the NMR experiment can also be calculated from the molecular dynamics simulation. ^64^ This strategy can serve as an interface with the experimental data to substantiate key findings on allostery. A protein graph connectivity network provides a deeper understanding of the information flow pathway inside the antibody. With the aid of graph theoretical measures like betweenness or eigenvector centrality, one can quantify the importance of individual resides in the allosteric information propagation. We specifically looked at the change in betweenness centrality for the antibody residues induced by the mutations in the spike antigen. While a group of amino acids in the B38 light chain (proximal to RBD residue 501) experience centrality reduction, it increases in virtually the opposite end of the heavy chain in the RBD-antibody interface. Absence of the N501Y mutation reduces this effect, revealing a antigen variant induced re-wiring mechanism that decreases the traffic of allosteric communication inside antibody in the vicinity of the N501Y mutation and increases near the K417N and E484K/Q mutations. This effect being particularly specific to the location of the mutation in the epitope, it explains the mutation specific collective change in the cross-correlation throughout the B38 antibody. In contrast, all the BD23 residues near the RBD binding interface experience a reduced centrality, irrespective of the specific location of the mutations. Some distant residues, following the trend of interfacial residues, show a change in BC value without the regard of the location of the mutation, explaining the specific RBD-BD23 cross-correlation pattern. These contradictory features point to inherent differences in the microscopic dynamics of the antibody, depending on the nature of mutations in the antigen and on the nature of the antibody. They have repercussions beyond the current pandemic, controlling the fundamental process of antigen-antibody recognition at a molecular level. Such allosteric communications can be present within the hACE2 receptor upon binding to the mutants of the coronavirus spike protein. But, because of the prime role of the glycan moieties in controlling the binding affinity between the spike and the receptor, the demonstration of such effects from inter-residue correlations can be more subtle.

Whether allosteric modulation of monoclonal antibodies can significantly change the effectiveness of the immune response still remains to be fully understood. Nevertheless, we demonstrated that allosteric perturbations do take place within the antibody and such perturbations can be connected to the mutations in the receptor-binding domain of the spike protein. To see if these mechanisms can be generalized to a wider range of pathogens and antibodies, an analysis similar to ours can be applied to other antigen-antibody pairs. We hope to verify this hypothesis in future work.

The process of unbinding of the antibodies from the spike proteins of different variants is also of significant interest, as it provides a more mechanistic insight into the mutation induced modification of the antibody binding propensity. Irrespective of the variant or the antibody, the RBD loop, consisting of residues 470-490, works like an anchor and is always the last region to interact with the antibody in the lowest energy dissociation pathways. The E484K/Q mutation in this region reduces the anchoring capability with the BD23 antibody by the loss of key hydrogen binding interactions. So, the N501Y, K417N-E484K-N501Y, and L452R-E484Q mutants show a decreasing trend in the non-equilibrium work required to dissociate the BD23 antibody, and the recovery of the E484 residue in the *delta* variant regains the binding affinity. B38, in contrast, does not interact with residues 484 or 452 in the bound state, and mutations in those residues have little to no impact on the interaction energy of B38 with the spike protein. So, all the RBD variants in this study have almost identical binding affinity with the B38 antibody, as apparent from the steered MD results. But the mutants have lower “binding strength” than the WT because of the loss of stabilizing hydrogen bonds as a result of the N501Y mutation. It is interesting to note that the newly emerging highly contagious *delta* (L452R-T478K), *delta plus* (K417N-L452R-T478K) and *omicron* ^65^ (S477N, T478K, E484A, Q493K etc.) variants also gained different mutations in this important loop region that is vital in preventing the dissociation of the neutralizing antibodies.

In summary, we quantified how the microscopic dynamics of the antigen-antibody complex is modulated by the mutations in the antigen and the nature of the antibody, with a particular focus on the SARS-CoV-2 spike protein. Although there is significant diversity in the dynamical perturbations observed at the molecular level, there are common underlying themes: modulation of inter-residue correlations, allosteric information propagation, and destabilization of non-bonded interactions dictate the stability of the antigen-antibody complex, which in turn determines the efficacy of monoclonal antibody therapies. Taken together, this study explores the molecular signatures of the antigen-induced allosteric effects within the antibodies for coronavirus. The scope of our findings is, however, broader and future studies in this direction can lead to a deeper understanding of the immune evasion mechanism of mutating viruses to help in designing adaptive therapeutic strategies to induce immune responses against new viral strains via monoclonal antibodies and vaccination.

## Methods

### System Preparation

The coordinates of the antibodies bound to the receptor binding domain (RBD) of the spike protein in wild type SARS-CoV-2, were obtained from the Protein Data Bank (PDB) with the following accession IDs: 7BZ5 (B38) ^31^ and 7BYR (BD23).^32^ Apart from the B38 antibody, the 7BZ5 PDB file contains the RBD domain (only residues 334-526) including one N-acetyl-*β*-D-glucosamine (NAG) moiety attached to Asn343. This glycan moiety was preserved in our calculation. The 7BYR PDB file includes the entire spike trimer along with the antibody, so we removed all spike residues except for 334-526 of the RDB bound to the antibody. The bare RBD systems were constructed from the 7BZ5 system by deleting the antibody. The mutations in the RBD were added using CHARMM-GUI input generator. ^66^ Each system was solvated in a cuboidal water box with 12 Å of padding in each direction, and Na^+^ and Cl*^−^* ions were added to neutralize the total charge and maintain an ionic strength of 150 mM. The hydrogen mass repartitioning (HMR) technique ^67^ was used to allow for 4 fs time-steps, facilitating longer simulation runs. The topology files for HMR were prepared using the VMD 1.9.4 alpha 51 software. ^68^ All systems were modeled using CHARMM36 force field^69^ (with CMAP correction for proteins) and TIP3P parameters^70^ were used for the water and the ions.

### Equilibration and Unbiased Simulation

Each system was minimized using a conjugate gradient algorithm for 10000 steps followed by an equilibration in NPT ensemble for 250 ps (with 2 fs time step) with harmonic restraint on the alpha carbons of all the protein residues. A 10 ns NPT equilibration was performed (with 4 fs time-step) afterwards for every system with no restraint applied on any atom. The temperature was controlled at 303.15 K using a Langevin thermostat with damping constant 1 ps*^−^*^1^, and a Langevin piston was used to control the pressure at 1 atm.

Production runs were performed for each RBD-antibody complex for 0.5 *µ*s and for each bare RBD system for 0.3 *µ*s. For all systems, (WT and four variants), the cumulative unbiased simulation time was 6.5 *µ*s. Only the last 0.4 *µ*s of each antibody bound system and 0.25 *µ*s of each bare RBD system were used for further analysis. All simulations were performed using NAMD 2.14 simulation package. ^71^ The trajectories were saved at an interval of 100 ps.

The root mean squared deviation (RMSD) and the residue-wise root mean squared fluctuations (RMSF) were computed using the gmx rms and gmx rmsf tools, respectively, in the GROMACS 2018.1 software. ^72^ RMSD for all systems were computed with respect to the first frame of the long production run. Trajectories were aligned only with respect to the *α*-carbons of the RBD using VMD software. ^68^ RMSD was computed for the RBD, antibody, the antigen binding domain (ABD) of the antibody and the whole system. The antigen binding domain is defined as the one of the two monomers of the BD23 protein which interacts with the RBD and the portions of the two monomers directly interacting with RBD (residue 1-108) for the B38 antibody (Fig. 1 in main text). For computing RMSF, the last 400 ns of each trajectory is split into 4 segments of 100 ns each. Residue wise RMSF was computed for each segment and the mean and standard deviations are reported. The RMSF ratio is then computed for each mutant strains as the ratio of RMSF for each residue and that of the corresponding residue in the WT. Principal component analysis was performed using the PyEMMA 2 package. ^73^ The non-bonding interaction energy (including the electrostatic energy and van-der Waals energy) between the antibody and the mutated residues of the RBD for different variants were calculated using the NAMD Energy module^74^ of VMD.^68^ Hydrogen bonds between the RBD and antibodies were also analyzed using VMD. Binding free energy between each RBD-antibody pair was computed using the Molecular Mechanics-Generalized Born Surface Area (MM-GBSA) approach ^75^ using the last 400 ns of the equilibrium trajectories. The NAMD2.14 package was used to perform the MM-GBSA calculation. ^76^

### Mutual Information-based Cross-Correlation and Protein Graph Connectivity Network

Linear mutual information (LMI) based cross-correlation matrices were computed for all residue pairs in each system. The theoretical details can be found in Refs. 77–79. Although the LMI-based cross-correlation is primarily used to quantify the allosteric effect of a small molecule ligand on the receptor protein, ^80, 81^ it can also be used to decipher the allosteric regulation within a protein or protein complex.^61^

The linear mutual information (LMI) metric (*I_ij_*) is calculated as

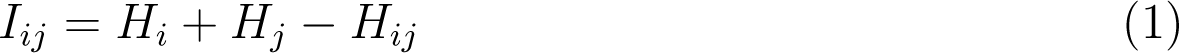

where *H* is a Shannon type entropy function:

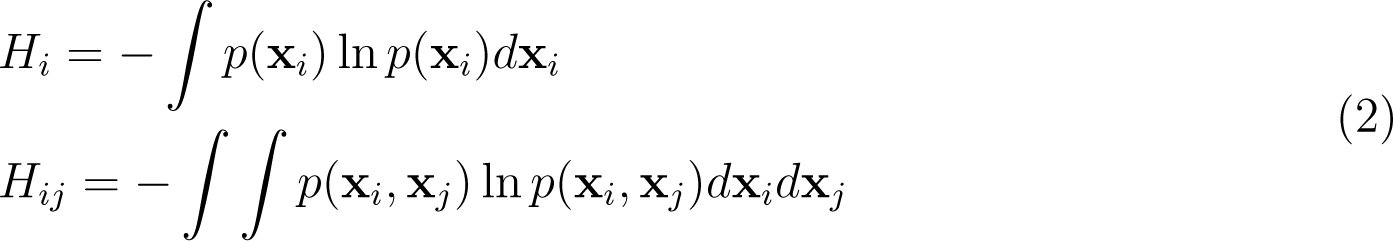

**x***_i_,* **x***_j_* are the vectors containing the Cartesian coordinates of the *C_α_* atoms, *p*(**x***_i_*) and *p*(**x***_j_*) are their marginal distributions and *p*(**x***_i_,* **x***_j_*) is their joint distribution. ^79, 81^ A Pearson like correlation coefficient *C_ij_* is defined from *I_ij_* in the following manner:

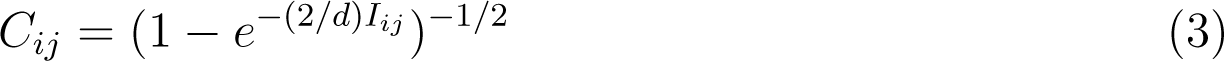

where *d* is the dimensionality of the space. Unlike the conventional correlation coefficient, *C_ij_* defined by Eq. can only take values between zero and one.

The residue-wise cross-correlation matrices were averaged over the four 100 ns segments prepared from the last 400 ns of each unbiased production run. The changes of the cross-correlation across the mutants were used to decipher the role of the mutations in the large scale dynamics. The uncertainty associated with the cross-correlations computed from the four 100 ns segments was found to be quite low, which allowed for relative comparison between different variants (Supporting Information Fig. S12). We also constructed a protein graph connectivity network model based on the cross-correlation matrices. The *α*-carbon of each residue is considered as the nodes of the graph and the cross-correlation between residues as the weight of the edges. Two residues were considered connected only if the heavy atoms of those residues were within 5 Å of each other during at least 75 % of the entire production run. The betweenness centrality (BC) of each residue in the network was computed and compared between various mutant strains. The BC quantifies the information flow between nodes or edges of a network. As the *α*-carbon of each residue is a node in the protein-graph-connectivity network, the value of the BC denotes the functional relevance of each residue in the dynamics of allosteric information flow through the protein chains. If a node, *i*, functions as a bridge between two other nodes along the shortest path joining those two, the BC of the node *i* is

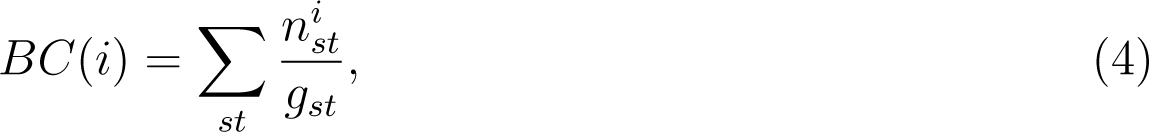

where *g_st_* is the total number of geodesics a.k.a. the shortest paths joining nodes *s* and *t*, out of which *n^i^* paths include the node *i*.

The change in BC, due to RBD mutations, is used to quantify the dynamical role of mutations in modifying the information flow pathway in the RBD-spike complex. The LMI based cross-correlations were calculated and the protein graph connectivity networks were constructed using the bio3D package. ^82^

### Steered Molecular Dynamics Simulation

To study the unbinding of the antibodies from the spike RBD, steered molecular dynamics (SMD) simulations^83^ were performed. For SMD simulation, all systems were modeled with the Generalized Born (GB) implicit solvent model^84^ with solvent accessible surface area (SASA) correction (GBSA). The ion concentration was set at 0.15 M. Instead of applying the pulling force to a specific atom or set of atoms, we defined a collective variable as the center of mass distance between the RBD and the antibody. A fictitious particle was attached to the pulling coordinate with a harmonic spring with a force constant of 1 kcal/mol/Å, to allow for sufficient fluctuations. This fictitious particle is pulled at a constant speed of 4 Å/ns. The external biasing potential applied to the system at time *t* can be given by:

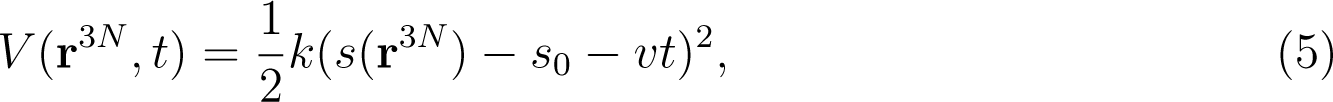

where *k* is the force constant of the harmonic spring, **r**^3^*^N^* is the 3*N* dimensional atomic coordinates, *s* is the collective variable, and *v* is the pulling speed. Five SMD trajectories of 10 ns length were generated for each system. Each trajectory was saved at an interval of 10 ps, and the pulling force and the accumulated work were recorded every 0.1 ps. The starting structures for the SMD trajectories were sampled from the unbiased production runs at the following points: 300 ns, 350 ns, 400 ns, 450 ns and 500 ns. All SMD simulations were started without further equilibration. The colvars^85^ module in conjunction with NAMD 2.14^71^ was used for SMD simulations. The SMD input files are provided in the GitHub repository and the SMD trajectories and the colvars output files are provided in the Zenodo repository (see Data Availability Statement).

We also performed additional SMD simulations in an explicit solvent environment to compare our results. The methodological details are provided in the supporting information.

## Acknowledgement

The authors thank Ly Le, Trevor Gokey, Sharon Stone, and Moises Romero for their suggestions and feedback on the manuscript. This work was supported partially by the National Science Foundation (NSF) via grant MCB 2028443. The authors thank the Triton Shared Computing Cluster (TSCC) in San Diego Supercomputer Center (SDSC), and University of California Irvine High Performance Computing (HPC) facility for providing the computational resources. The authors declare no competing financial interest.

## Data Availability Statement

All the trajectory data are available from the Zenodo Repository: 5281/zenodo.5676393. The codes and analysis scripts are available from the GitHub repository:

## Author Contributions

DR designed research with inputs from IA. DR and RNQ performed unbiased molecular dynamics simulations and analyzed results. DR performed SMD simulations and analyzed results. DR constructed the protein-graph connectivity network model using mutual information based cross correlation. DR and IA wrote the paper. All authors contributed to discussing the results and reviewing the manuscript.

## Supporting Information Available

Table S1: List of all simulations performed in this study Protocol and results for the steered MD simulation in explicit solvent.

Figure S1-S27: Results of RMSD, RMSF, PCA, mutual information, betweenness centrality, hydrogen-bonding analysis, and interaction energy measurements of the RBD-Antibody systems and RBD only systems. Results of Steered MD simulation and the antibody dissociation mechanism for all variants.

## Supporting information

**Table 1:** A list of simulations performed in the current study

| System | Variant | Strain | RBD<br>Mutations | Unbiased<sup>a</sup><br>MD | Number<sup>a</sup><br>of Atoms | SMD<sup>b</sup> | Number<sup>b</sup><br>of Atoms |
| --- | --- | --- | --- | --- | --- | --- | --- |
| RBD only | WT | WT | - | $0.3\mu\text{s}\times 1$ | 31392 | - | - |
| | alpha | B.1.1.7 | N501Y | $0.3\mu\text{s}\times 1$ | 31477 | - | - |
| | beta | B.1.351 | N501Y,K417N,E484K | $0.3\mu\text{s}\times 1$ | 31384 | - | - |
| | kappa | B.1.617 | L452R,E484Q | $0.3\mu\text{s}\times 1$ | 31165 | - | - |
| | delta | B.1.617.2 | L452R,T478K | $0.3\mu\text{s}\times 1$ | 31716 | - | - |
| RBD<br>+<br>B38 Ab | WT | WT | - | $0.5\mu\text{s}\times 1$ | 76753 | $10\text{ns}\times 5$ | 9513 |
| | alpha | B.1.1.7 | N501Y | $0.5\mu\text{s}\times 1$ | 76916 | $10\text{ns}\times 5$ | 9520 |
| | beta | B.1.351 | N501Y,K417N,E484K | $0.5\mu\text{s}\times 1$ | 77500 | $10\text{ns}\times 5$ | 9519 |
| | kappa | B.1.617 | L452R,E484Q | $0.5\mu\text{s}\times 1$ | 75110 | $10\text{ns}\times 5$ | 9526 |
| | delta | B.1.617.2 | L452R,T478K | $0.5\mu\text{s}\times 1$ | 80889 | $10\text{ns}\times 5$ | 9532 |
| RBD<br>+<br>BD23 Ab | WT | WT | - | $0.5\mu\text{s}\times 1$ | 52625 | $10\text{ns}\times 5$ | 6491 |
| | alpha | B.1.1.7 | N501Y | $0.5\mu\text{s}\times 1$ | 50291 | $10\text{ns}\times 5$ | 6498 |
| | beta | B.1.351 | N501Y,K417N,E484K | $0.5\mu\text{s}\times 1$ | 53669 | $10\text{ns}\times 5$ | 6497 |
| | kappa | B.1.617 | L452R,E484Q | $0.5\mu\text{s}\times 1$ | 53782 | $10\text{ns}\times 5$ | 6504 |
| | delta | B.1.617.2 | L452R,T478K | $0.5\mu\text{s}\times 1$ | 55535 | $10\text{ns}\times 5$ | 6510 |

<sup>a</sup> Performed in explicit solvent molecules and ions.
<sup>b</sup> Performed in implicit solvent environment

## Steered Molecular Dynamics in Explicit Solvent

### Methods

To establish the validity of our implicit solvent SMD approach, we performed additional steered molecular dynamics simulations in an explicit solvent environment. The solvated structures were prepared following a protocol similar to that used for unbiased MD simulations, except we added 50 Å of water padding in one side along the principal axis of the RBD-Antibody complex, to allow for sufficient solvent coverage for the dissociating system. The pulling speed was kept at 1 Å/ns along the center of mass distance based CV to allow for sufficient time for the relaxation of explicit water molecules. This protocol led to a simulation time pf 40 ns for each complex. As these simulations are significantly more expensive compared to the implicit solvent SMD runs, only one set of pulling simulation was conducted for the WT and the *kappa* variant for each antibody. The choice of the *kappa* variant was motivated by the fact that in implicit solvent environment, this variant showed the highest change in the non-equilibrium work needed for detachment from the antibody in comparison to the WT.

### Results

The mutations in the *kappa* variant reduces the non-equilibrium work required for the dissociation of both the BD23 and B38 antibody (Fig. S26). This observation is consistent with our findings from the implicit solvent SMD simulations. Due to the presence of explicit water and a lower pulling force, we cannot expect the absolute value of the work to be identical. The dissociation mechanism of the RBD-antibody complex obtained from the implicit solvent and explicit solvent SMD simulation are in qualitative agreement with each other. In both cases, the RBD loop consisting of residues 470-490 works like an anchor to hold on to the antibody, immediately prior to the detachment (Fig. S27). The S477 residue of WT RBD and the A475 residue of the *kappa* variant RBD forms the last hydrogen bond with the B38 antibody during the force induced dissociation process, in both explicit and implicit solvent simulations. Although, the E484 of WT RBD form the last H-bond with the departing BD23 antibody under both conditions, the last connecting residue is different for the *kappa* RBD in the explicit (N481) and implicit (S477) environment. Although this does not change our overall conclusion that the mutations in RBD significantly perturbs the dissociation mechanism because the 484 residue which form hydrogen bond with the BD23 antibody looses its capability to form the hydrogen bond when it is mutated.

## Supporting Figures

**Figure S1:**
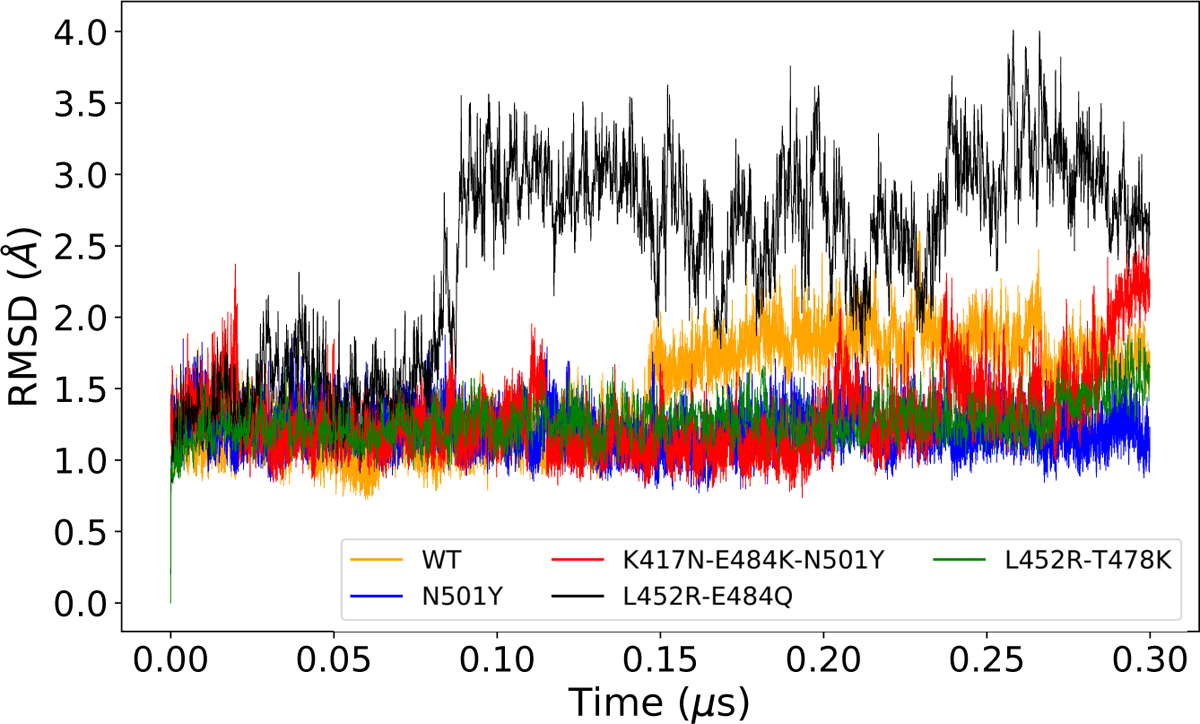
Root mean squared deviations of the bare RBD systems (WT and all mutants) from the long unbiased MD simulation.

**Figure S2:**
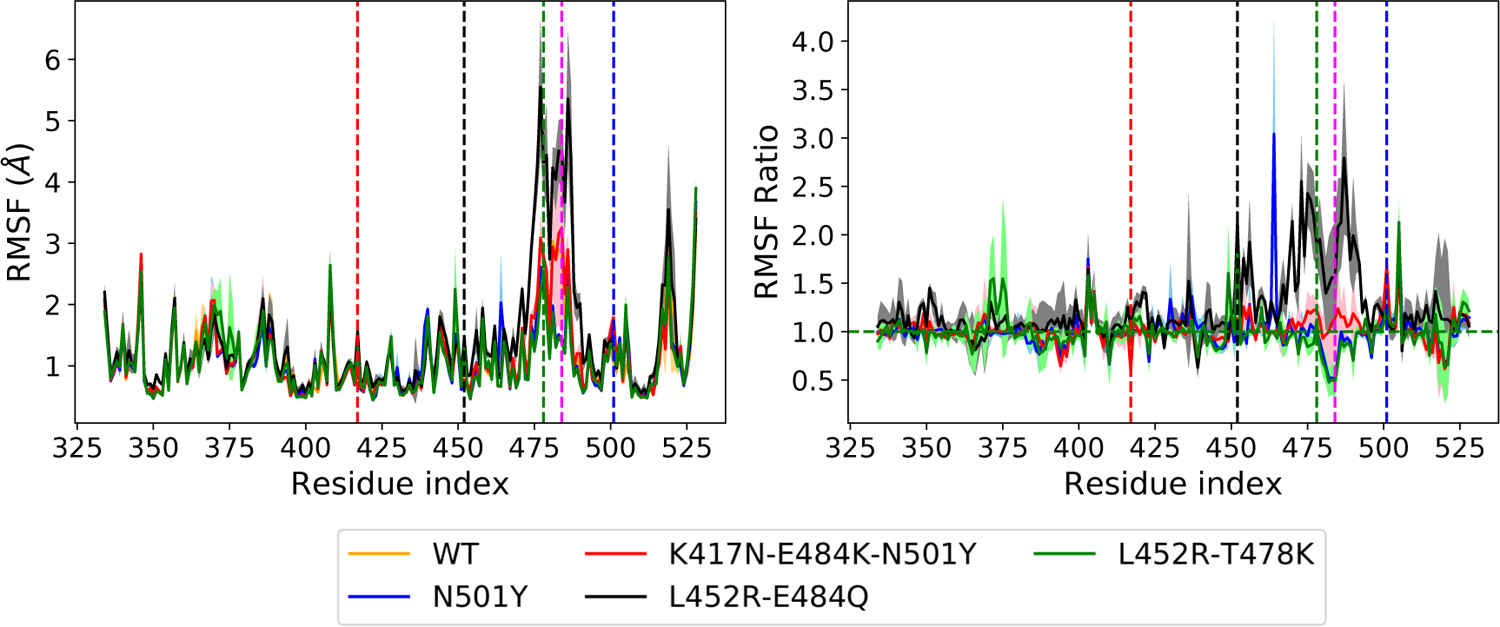
The root mean squared fluctuations (RMSF) of all residues of the bare RBD systems (WT and mutants) and the residue wise RMSF ratio for the three mutant systems.

**Figure S3:**
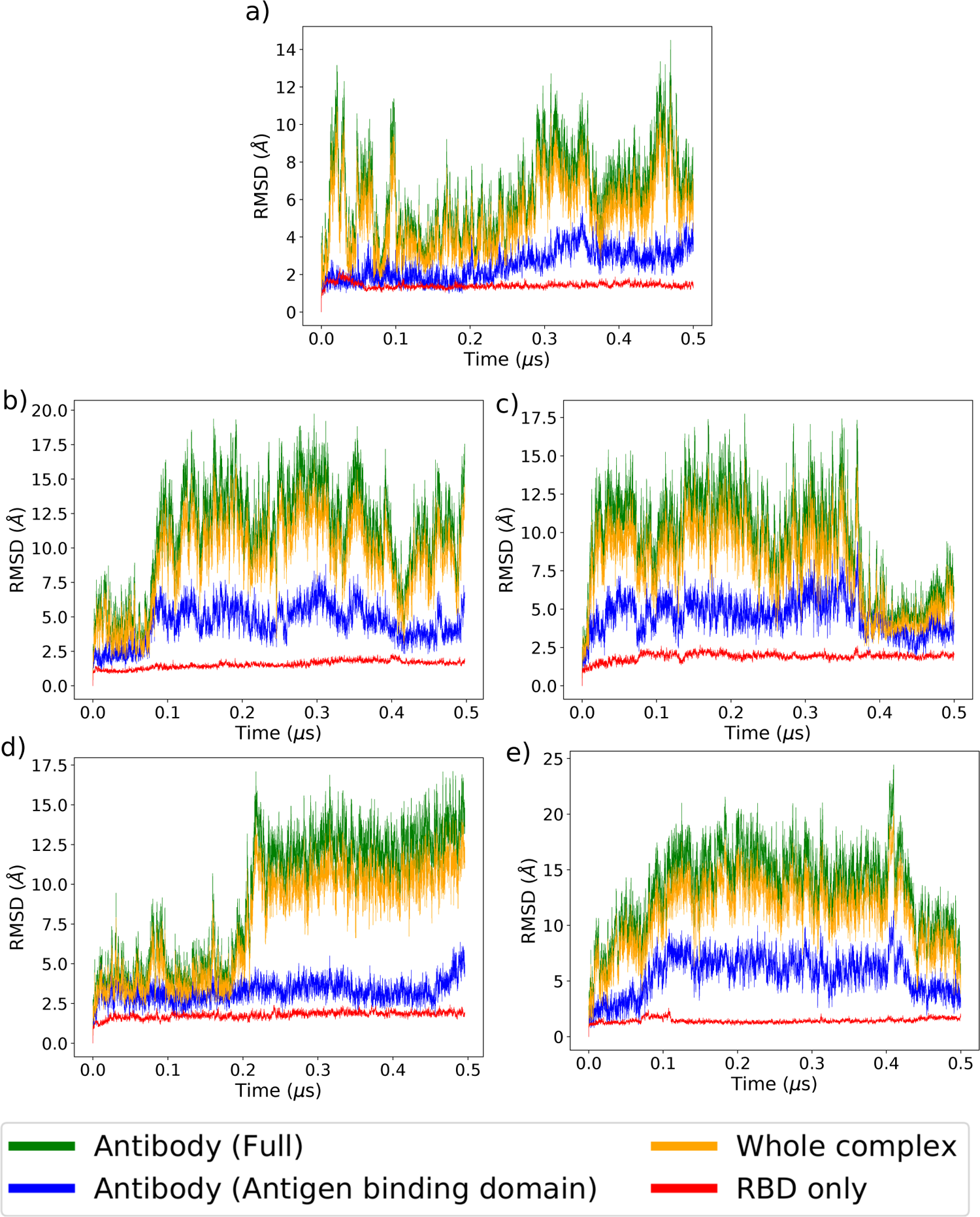
RMSD of different domains of the B38 Fab complex with (a) WT, (b) N501Y mutant, (c) K417N-E484K-N501Y mutant, (d) L452R-E484Q, and (e) L452R-T478K mutant forms of RBD from the long unbiased MD simulation.

**Figure S4:**
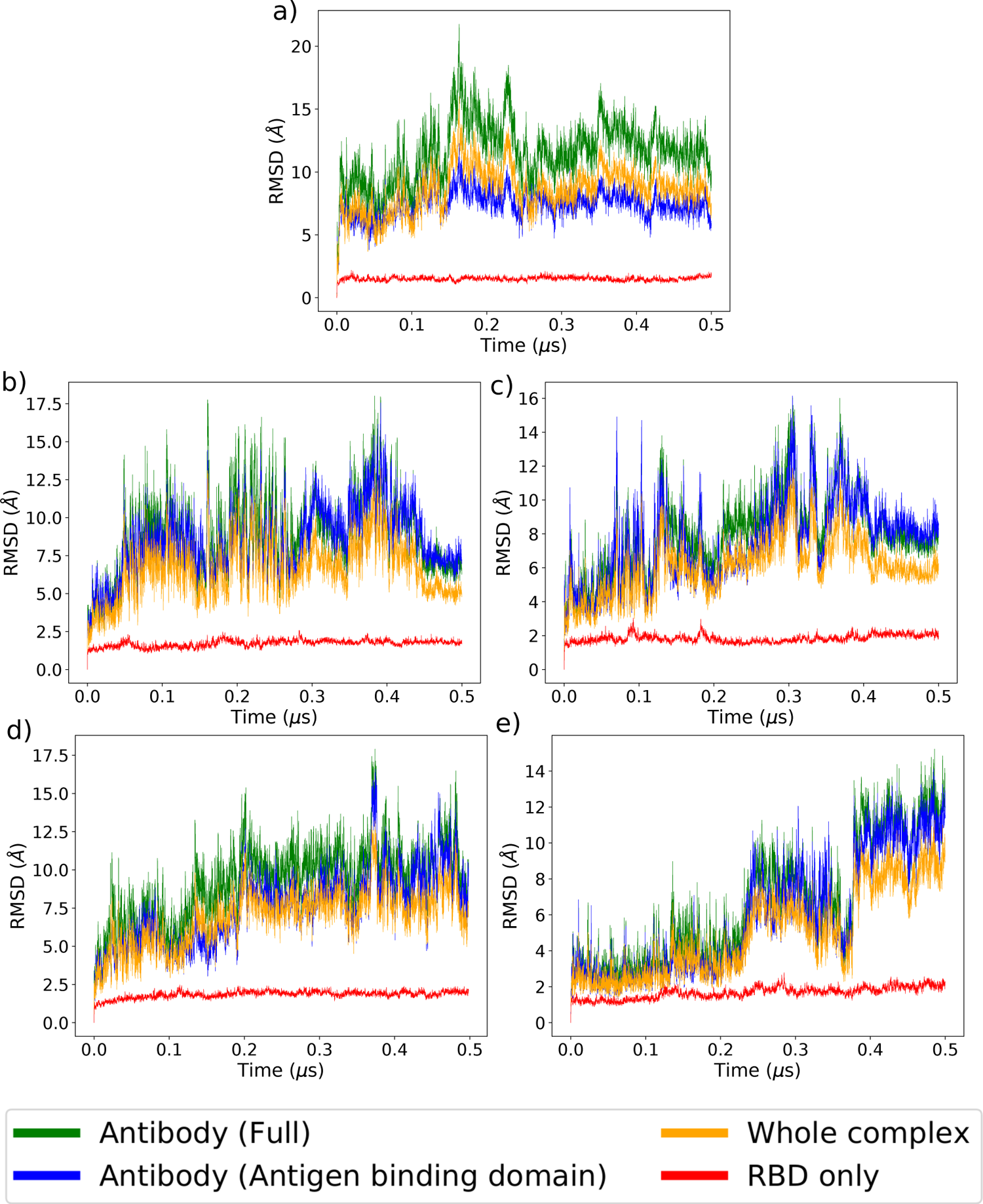
RMSD of different domains of the BD23 Fab complex with (a) WT, (b) N501Y mutant, (c) K417N-E484K-N501Y mutant, (d) L452R-E484Q, and (e) L452R-T478K mutant forms of RBD from the long unbiased MD simulation.

**Figure S5:**
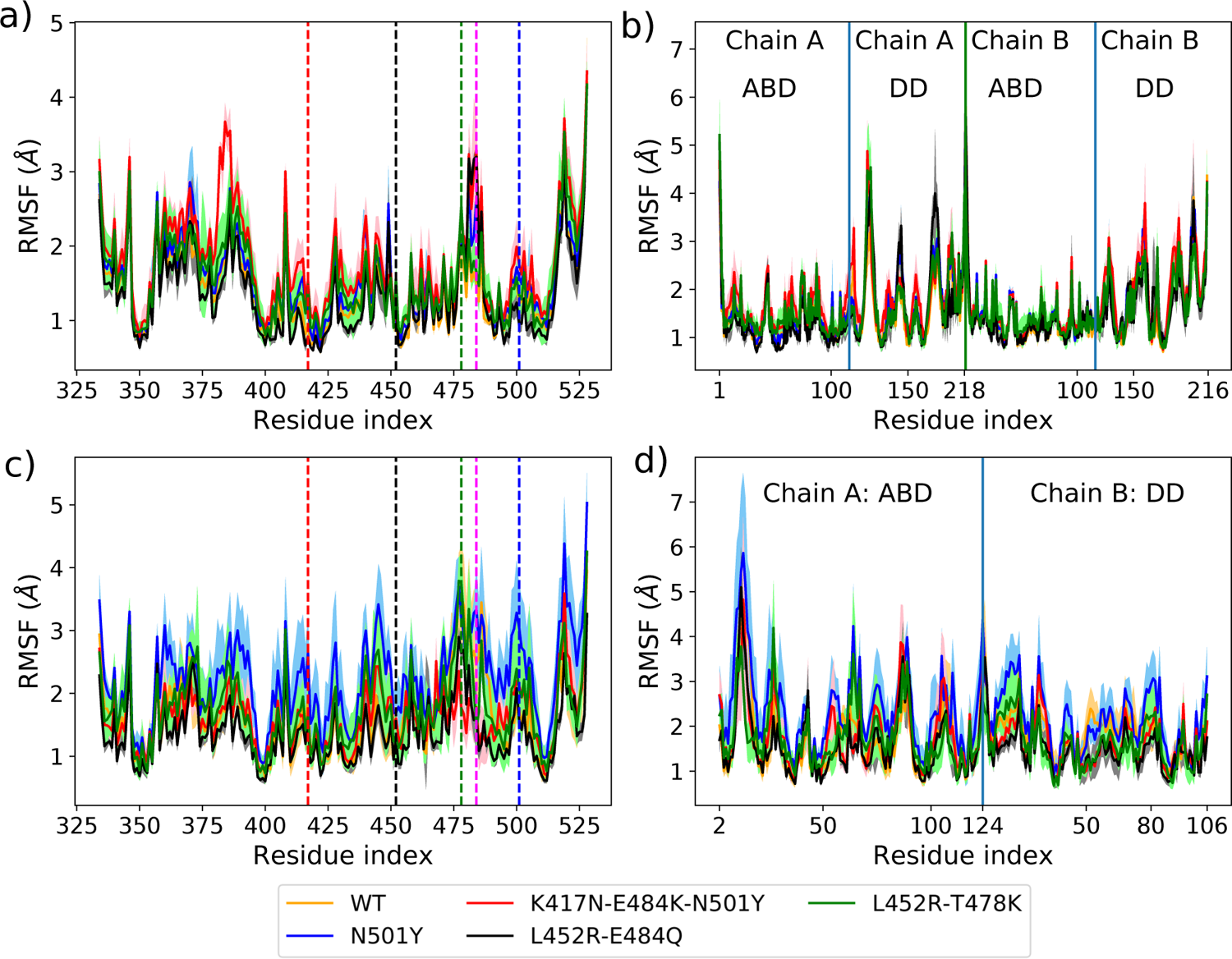
Residue-wise RMSF of all the RBD-antibody complexes: (a) RBD and (b) the antibody part of the RBD-B38 Fab complex and the (c) RBD and (d) the antibody part of the RBD-BD23 Fab complex, for the WT and the mutants. In (a) and (c) the location of mutated residues are marked in dashed line with the following color code; red: 417, black: 452, green: 478, pink: 484 and blue: 501.

**Figure S6:**
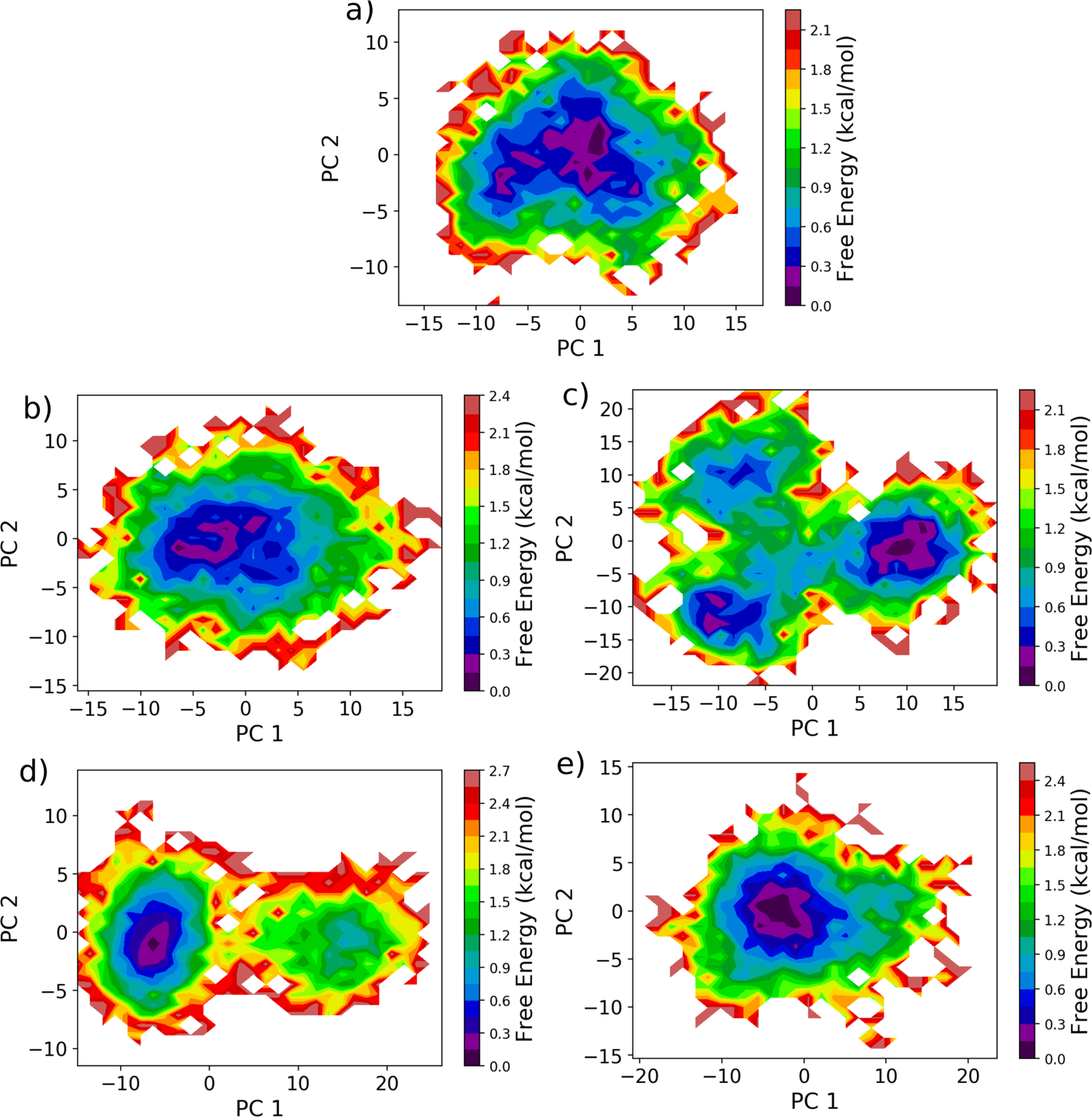
The free energy landscape of the RBD-B38 Fab complexes projected along the first two principal components, for (a) the WT, (b) N501Y mutant, (c)K417N-E484K-N501Y mutant, (d) L452R-E484Q, and (e) L452R-T478K mutant forms.

**Figure S7:**
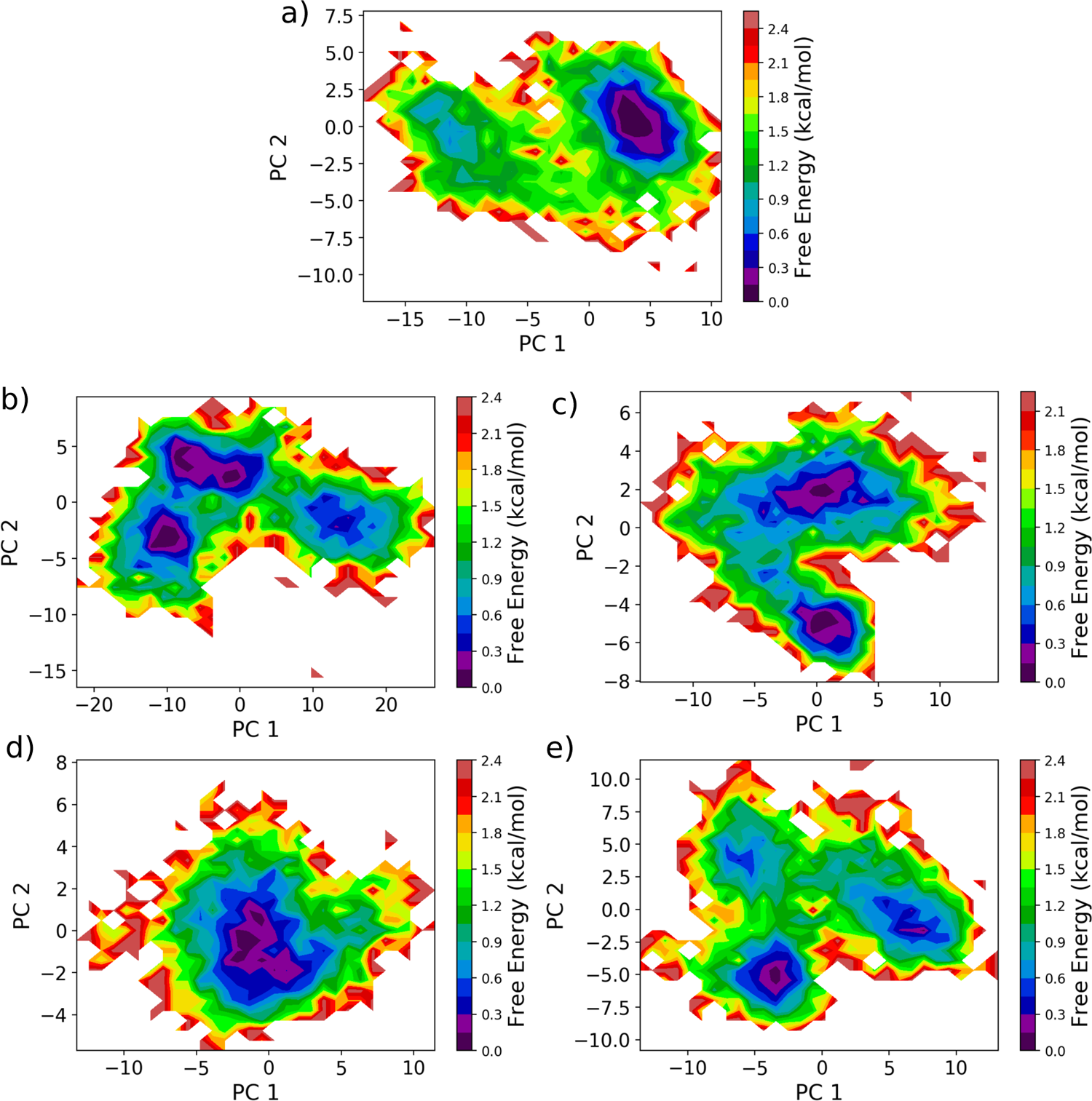
The free energy landscape of the RBD-BD23 Fab complexes projected along the first two principal components, for (a) the WT, (b) N501Y mutant, (c)K417N-E484K-N501Y mutant, (d) L452R-E484Q, and (e) L452R-T478K mutant forms.

**Figure S8:**
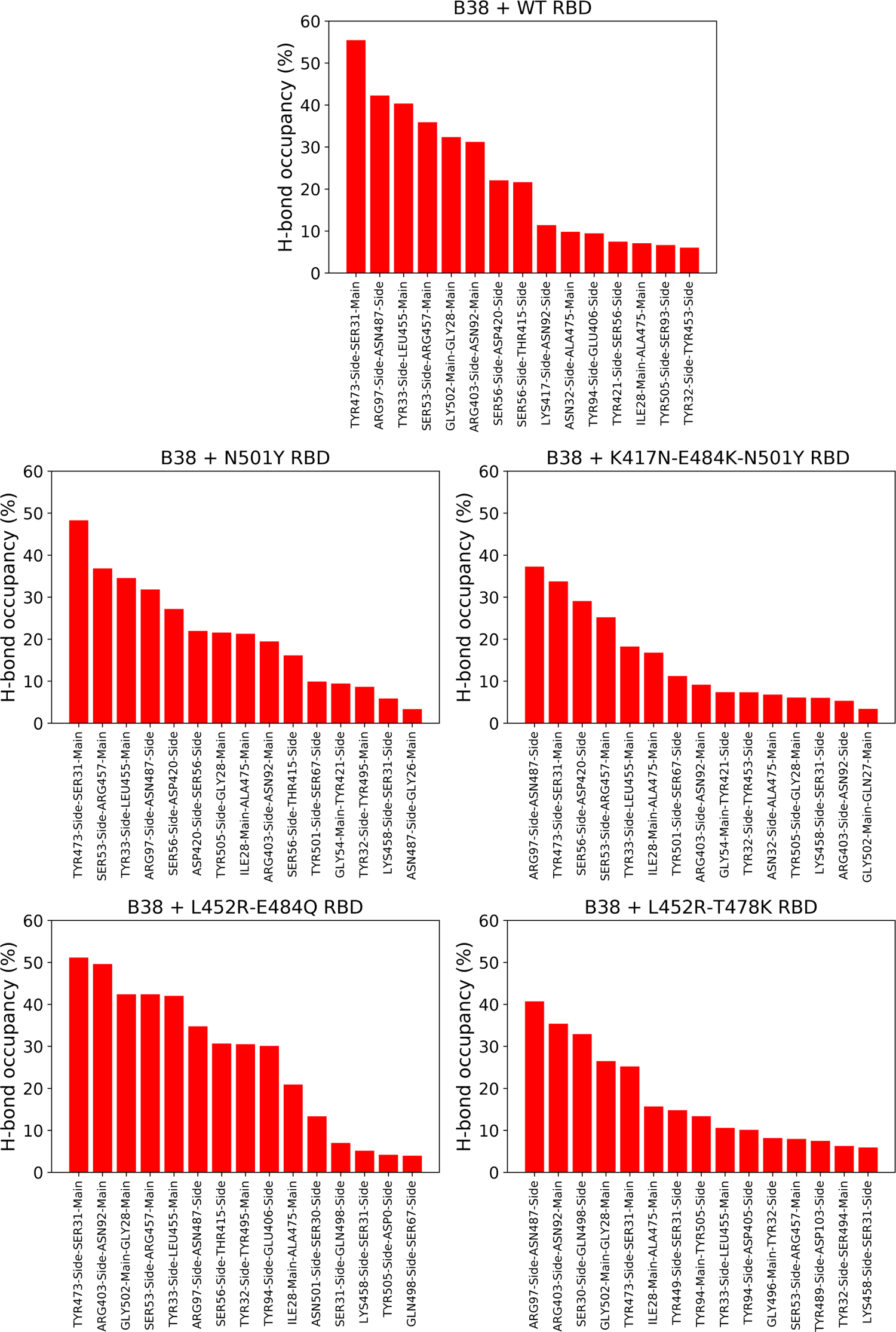
The occupancy of the most prevalent hydrogen bonds between the different variants of the spike RBD and the B38 antibody, sampled from long unbiased simulation.

**Figure S9:**
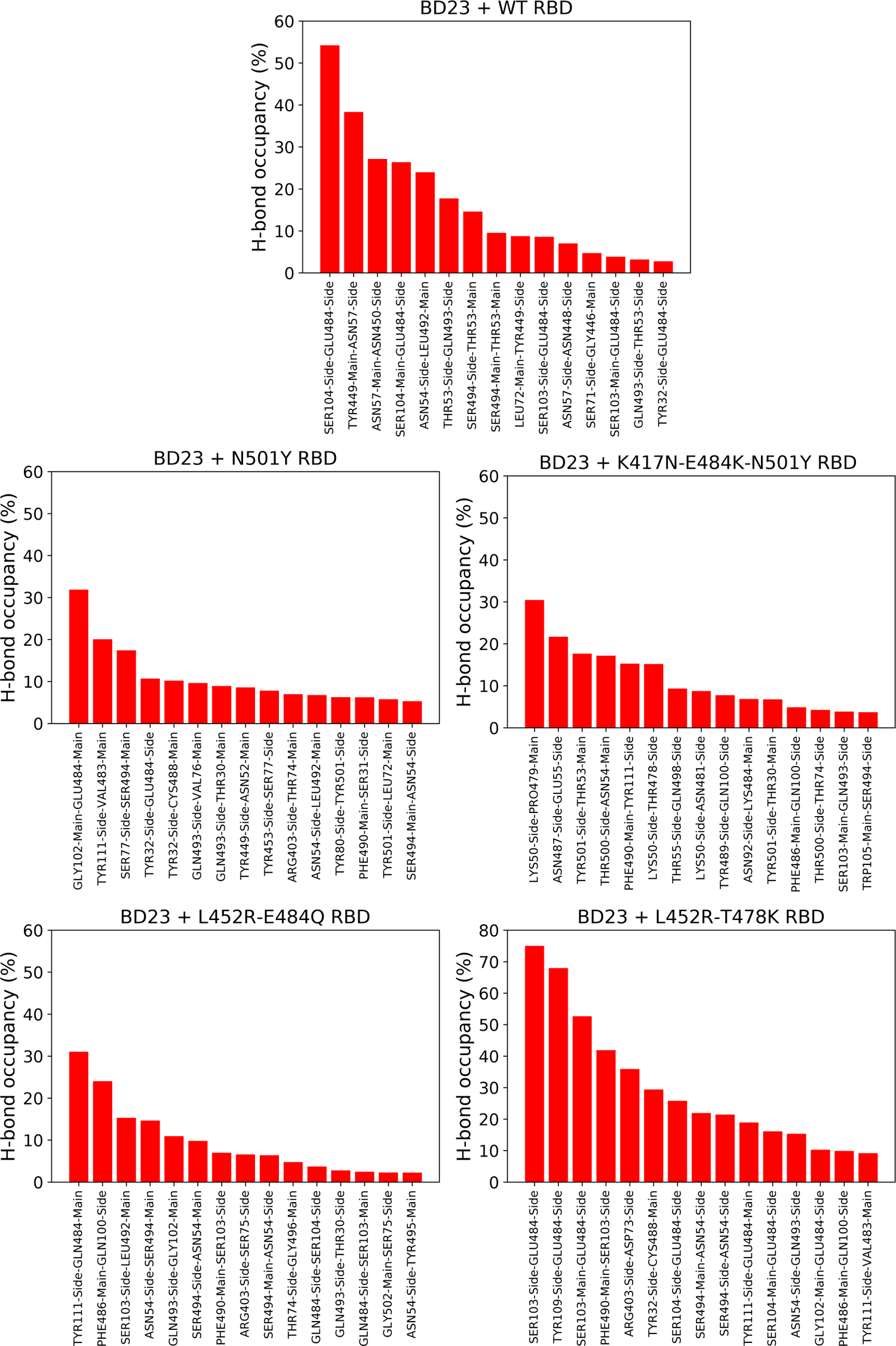
The occupancy of the most prevalent hydrogen bonds between the different variants of the spike RBD and the BD23 antibody, sampled from long unbiased simulation.

**Figure S10:**
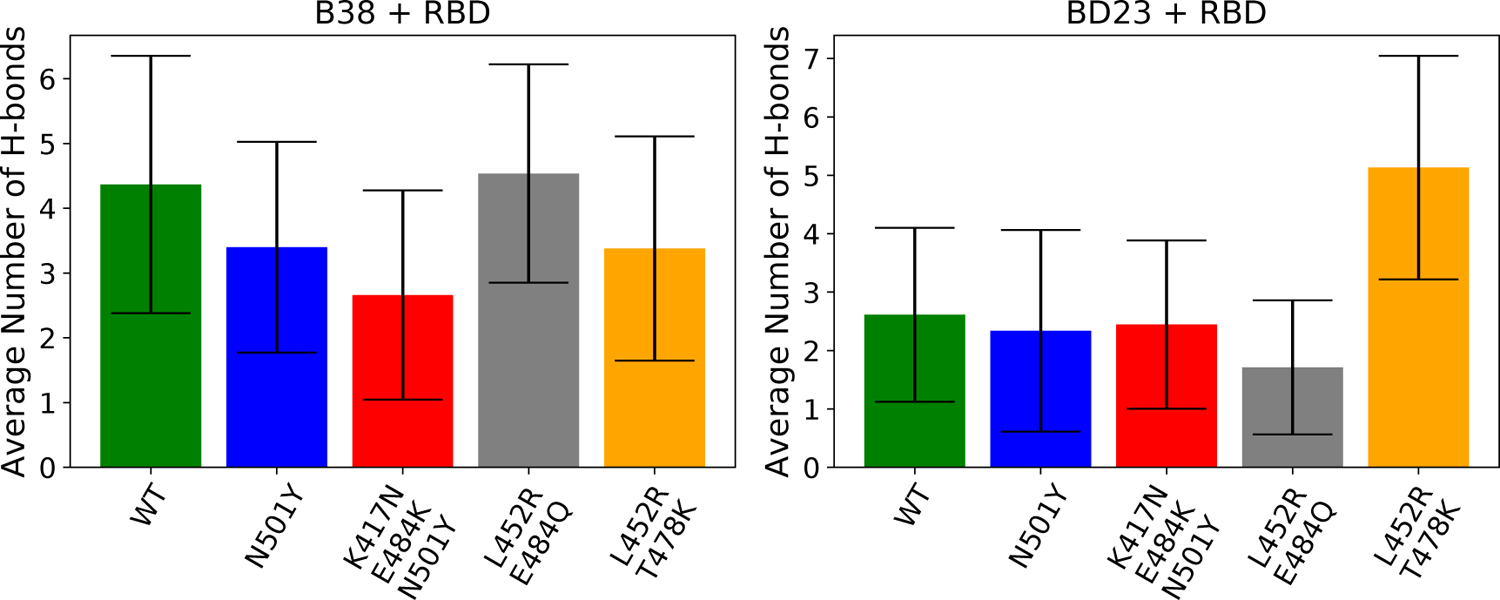
The average number of hydrogen bonds between different variants of the spike RBD and the two antibodies.

**Figure S11:**
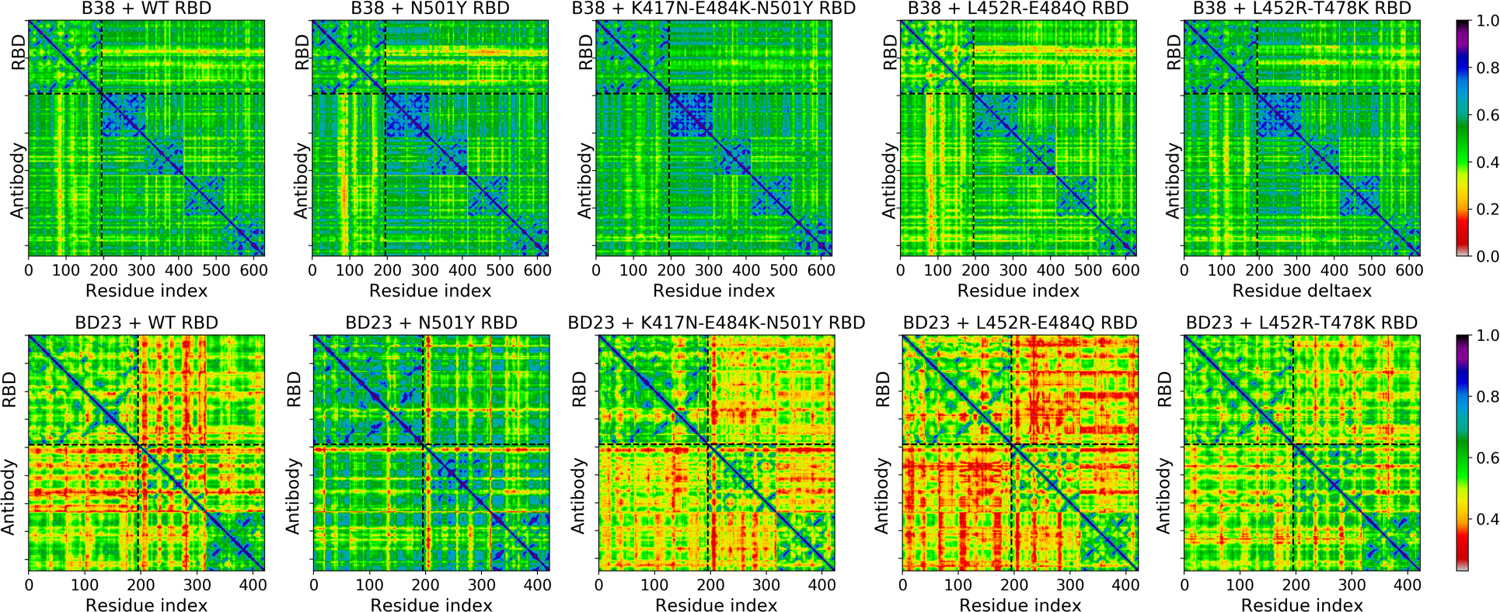
Linear mutual information based cross-correlation between all residue pairs in the RBD-antibody complexes for both the antibodies and all four variants of the RBD.

**Figure S12:**
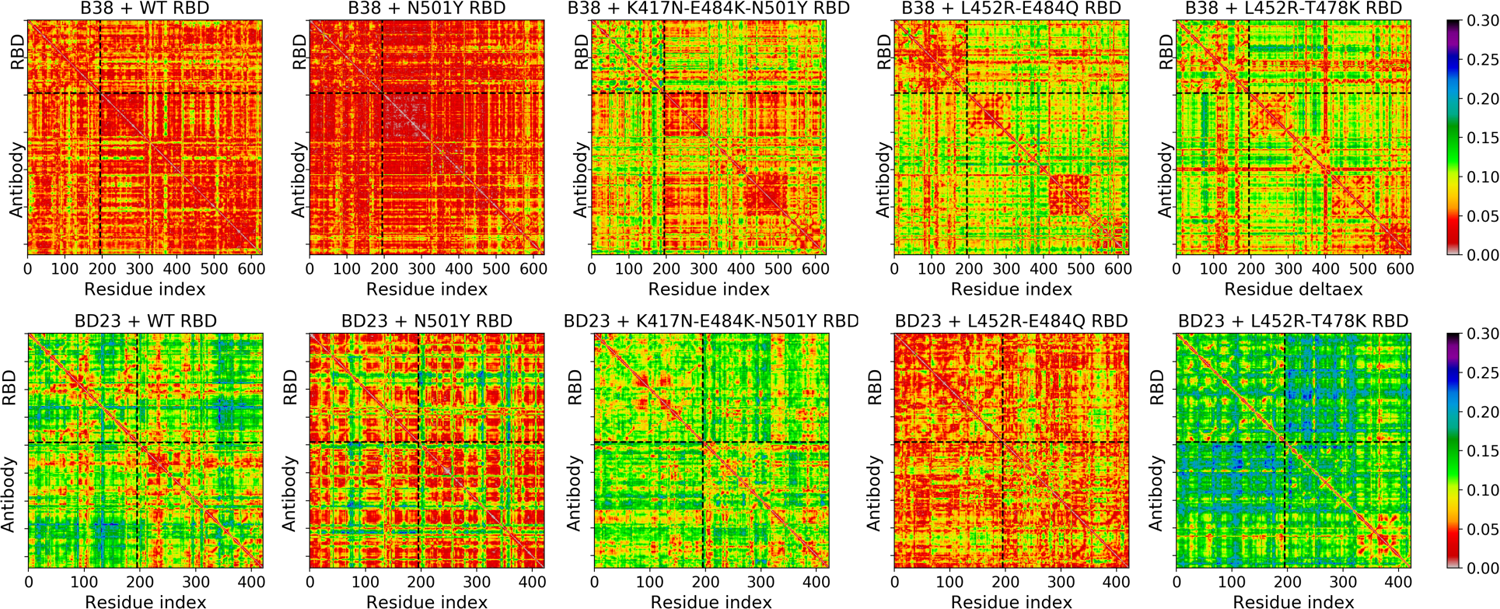
The standard deviation of the linear mutual information based cross-correlation between all residue pairs in the RBD-antibody complexes for both the antibodies and all four variants of the RBD. The standard deviation has been computed over four 100 ns trajectory segments (see Methods section in main manuscript)

**Figure S13:**
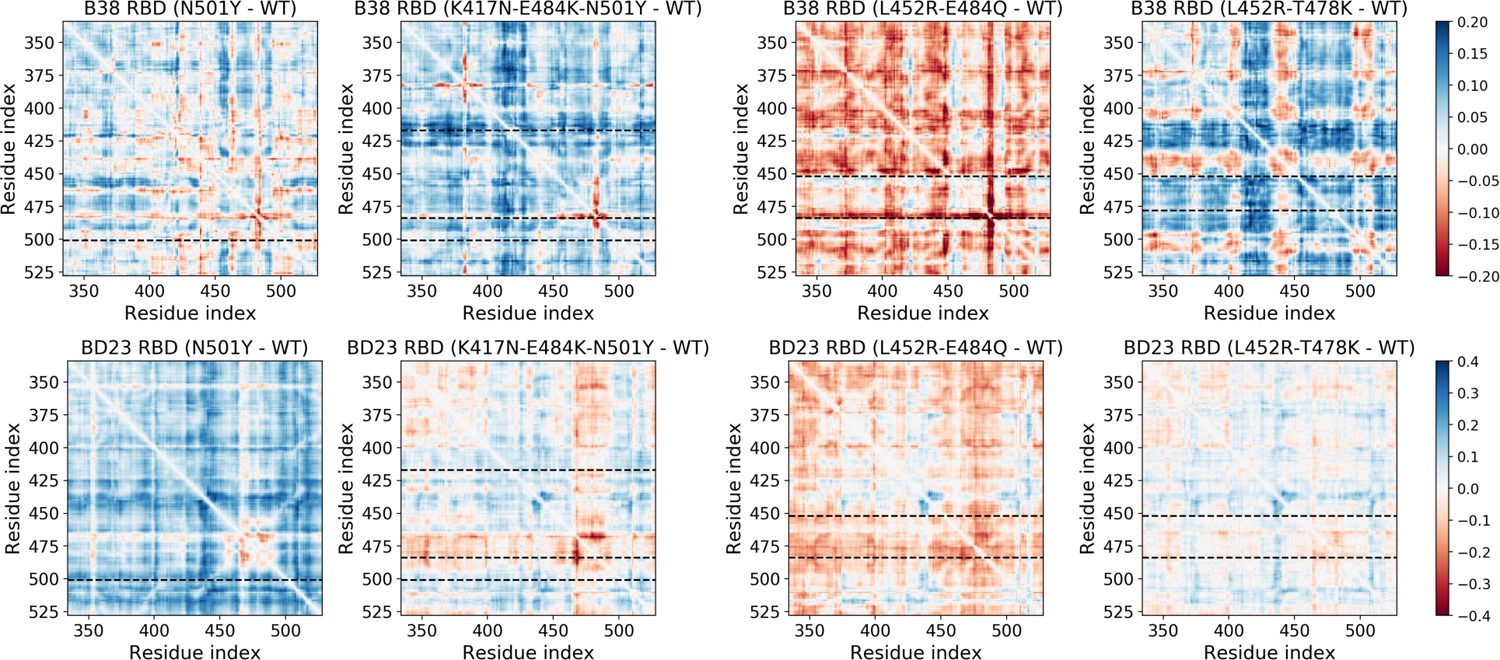
The change in the linear mutual information based cross-correlation between all residue pairs in only the RBD region for different variants and for two antibody systems. The WT-RBD system is considered as reference.

**Figure S14:**
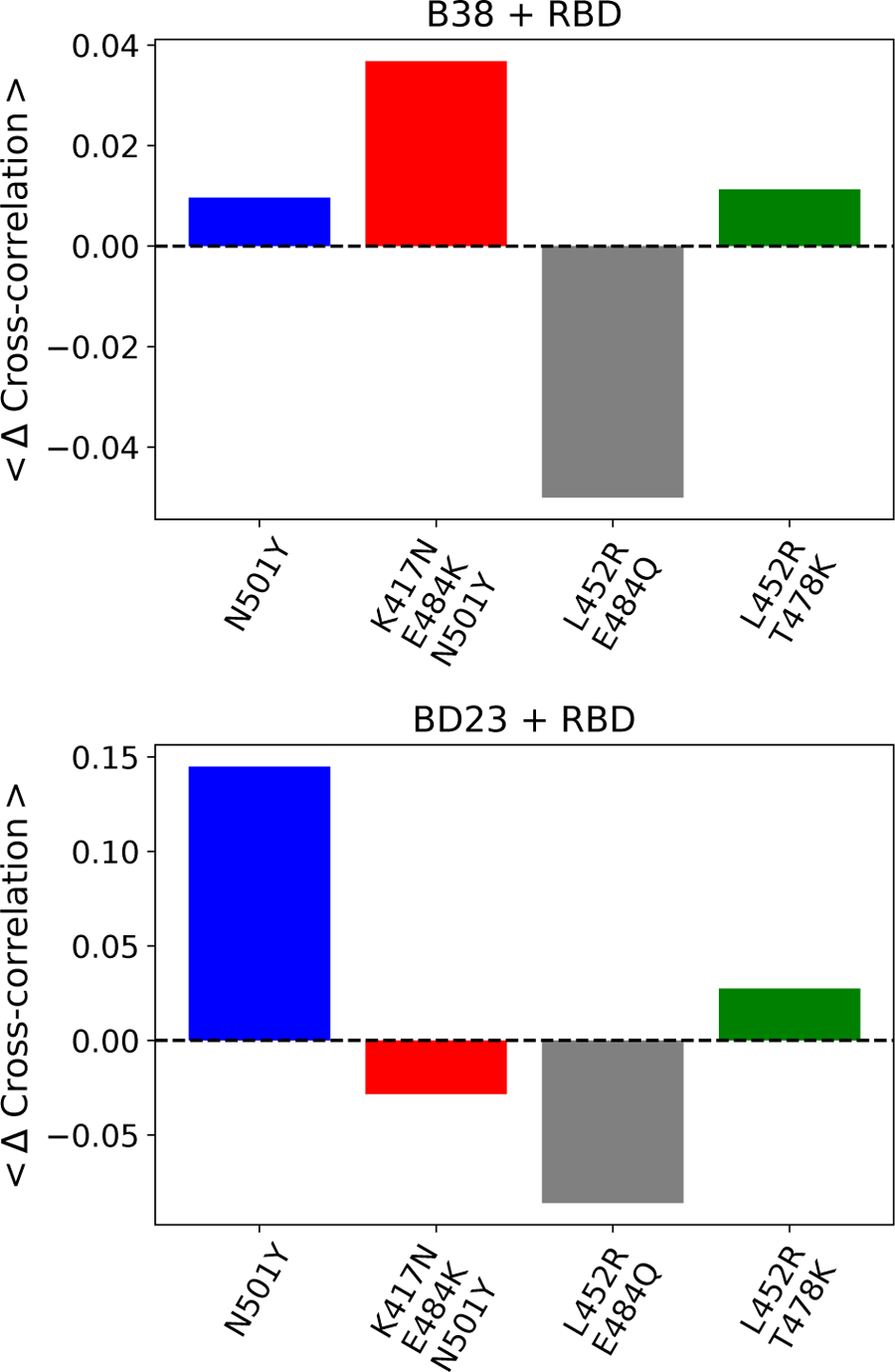
The average change in cross-correlation values per residue for the four mutants in complex with the B38 and BD23 antibody.

**Figure S15:**
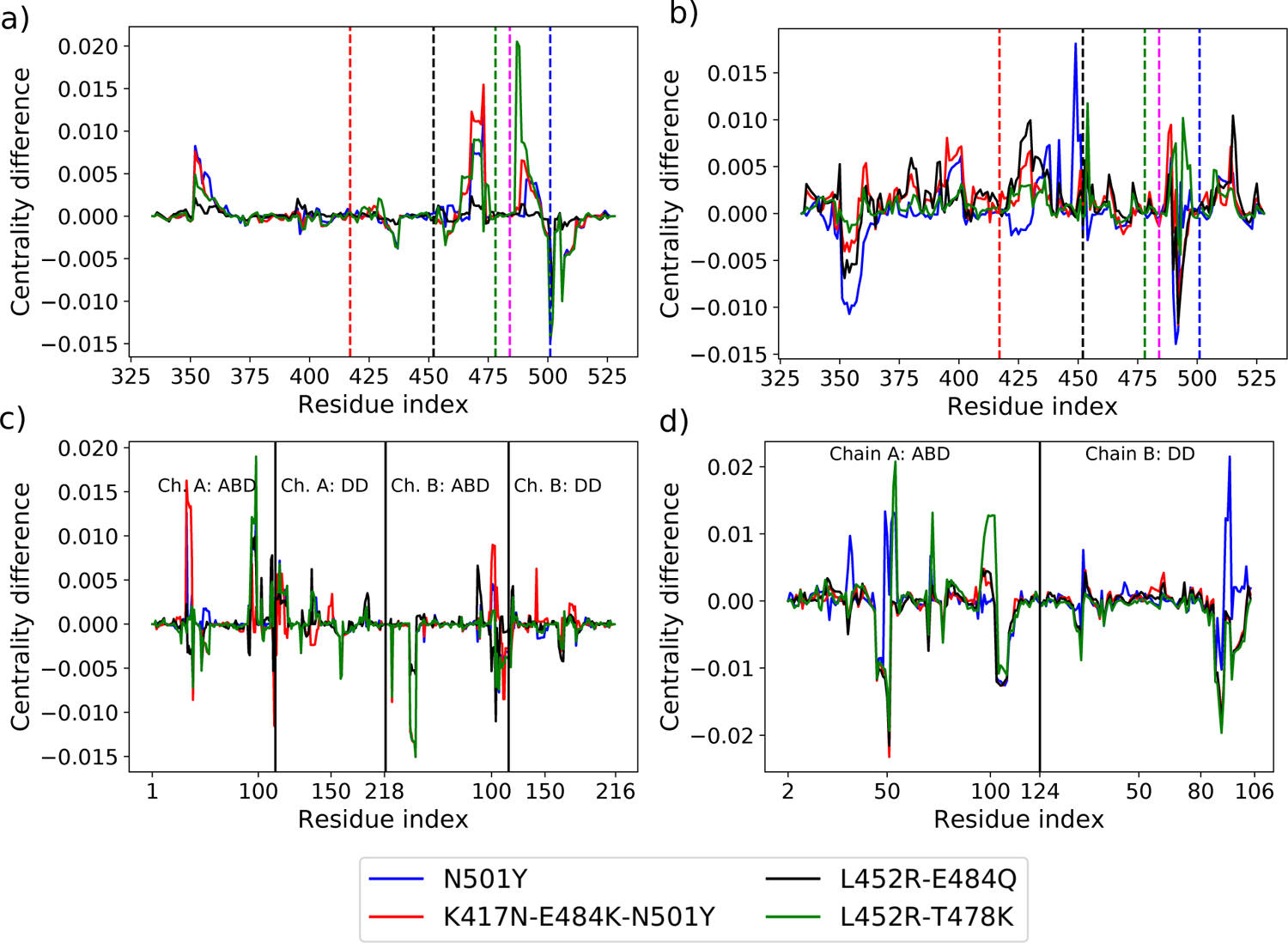
The change in betweenness centrality for all the residues of the mutant RBD complexed with the two antibodies. The upper panel depicts the BC change for only the RBD part for the (a) B38 complex and (b) BD23 complex. The location of the mutations are shown as vertical dashed lines. The lower panel depicts the same for only the antibody part for the (a) B38 complex and (b) BD23 complex.

**Figure S16:**
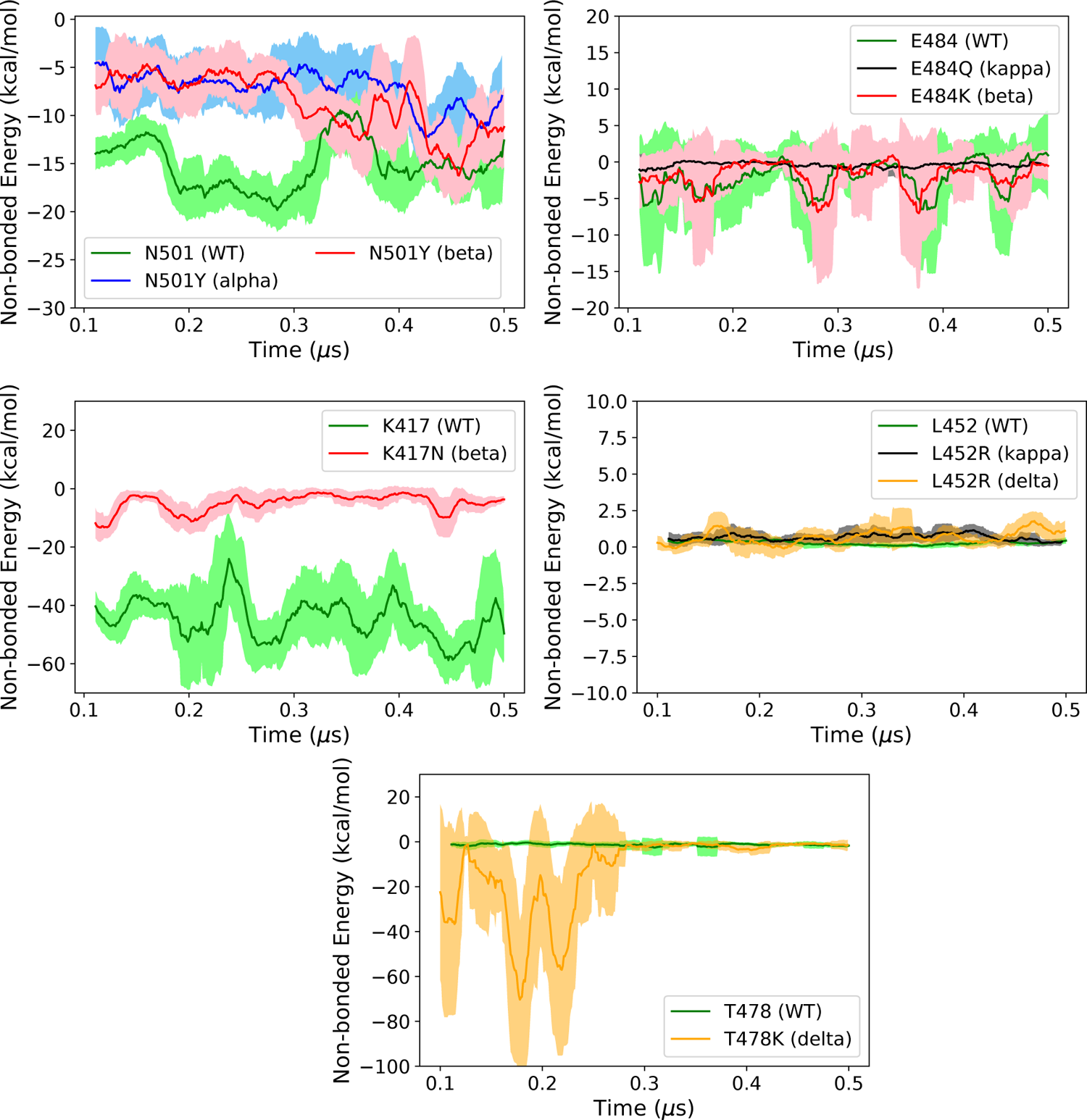
The non-bonded interaction energy between the B38 antibody and each individual RBD residue, mutated in the different variants. The error bars are running average over 20 ns.

**Figure S17:**
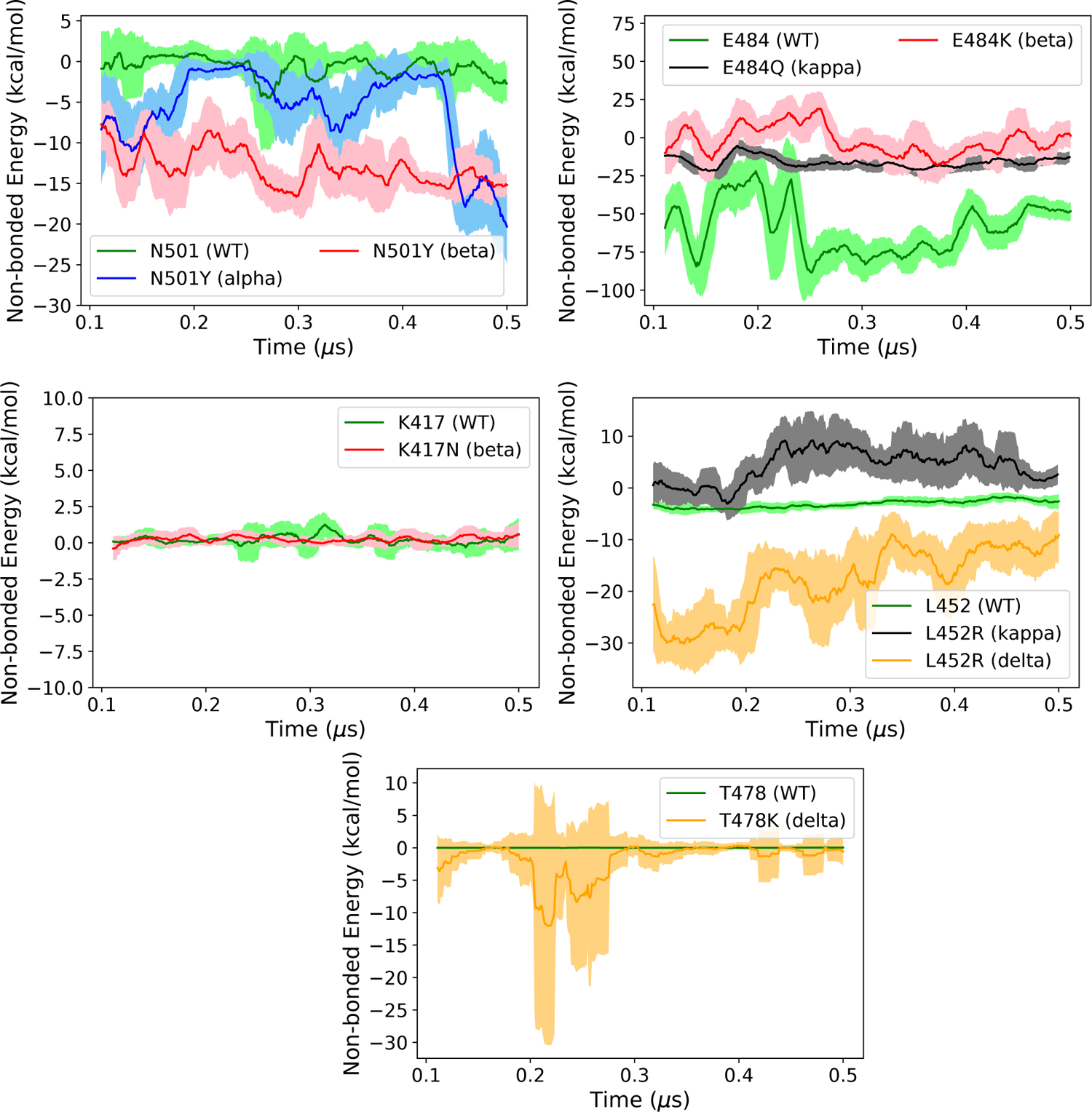
The non-bonded interaction energy between the BD23 antibody and each individual RBD residue, mutated in the different variants. The error bars are running average over 20 ns.

**Figure S18:**
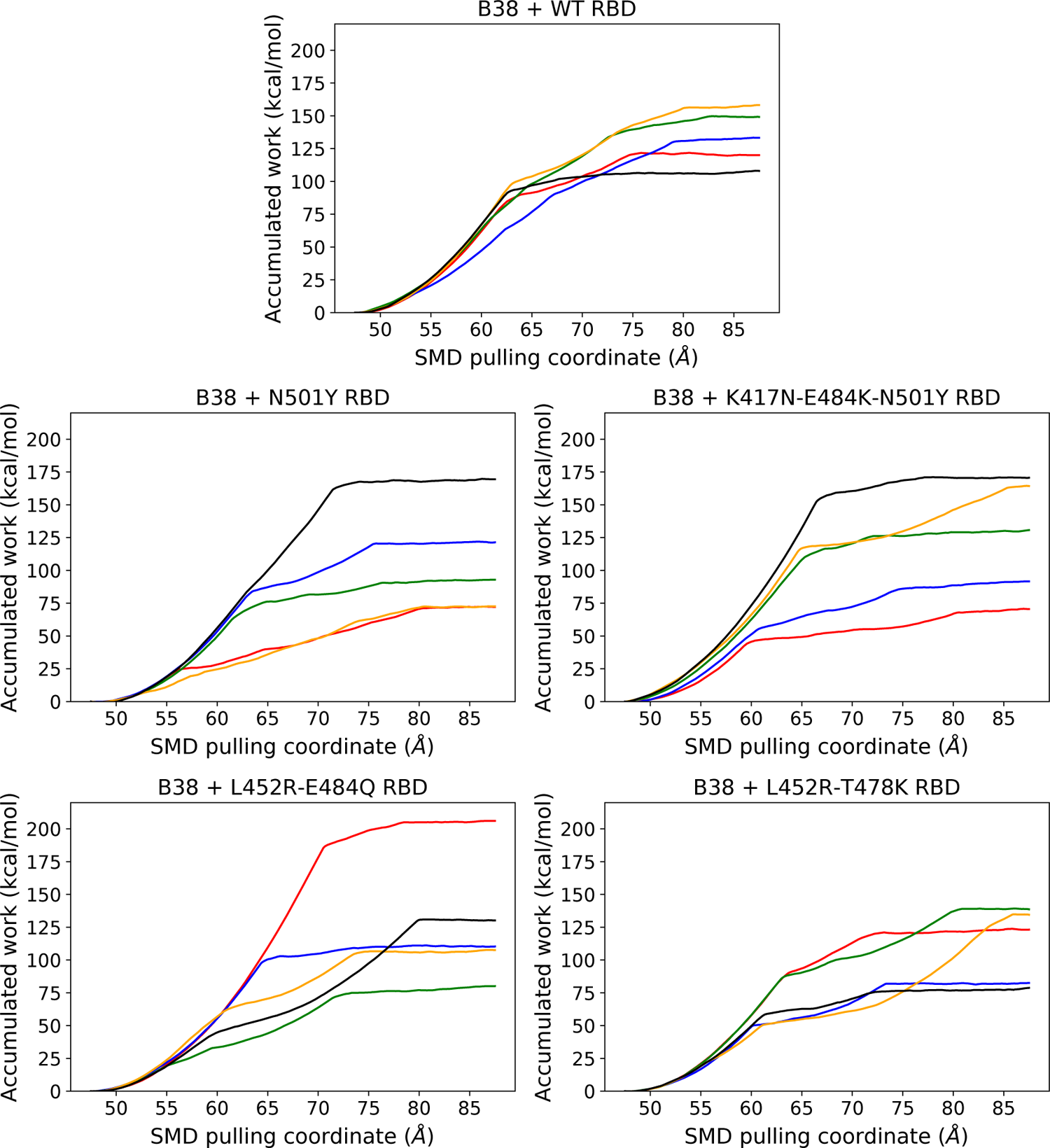
The work required to pull the B38 antibody along the center of mass distance between the RBD and antibody in the SMD simulation. The starting structures for each SMD simulation is sampled from long unbiased simulation after 300, 350, 400, 450 and 500 ns. The color scheme for these five SMD trajectories are as follows: red: 300 ns, blue: 350 ns, green: 400 ns, orange: 450 ns, and black: 500 ns.

**Figure S19:**
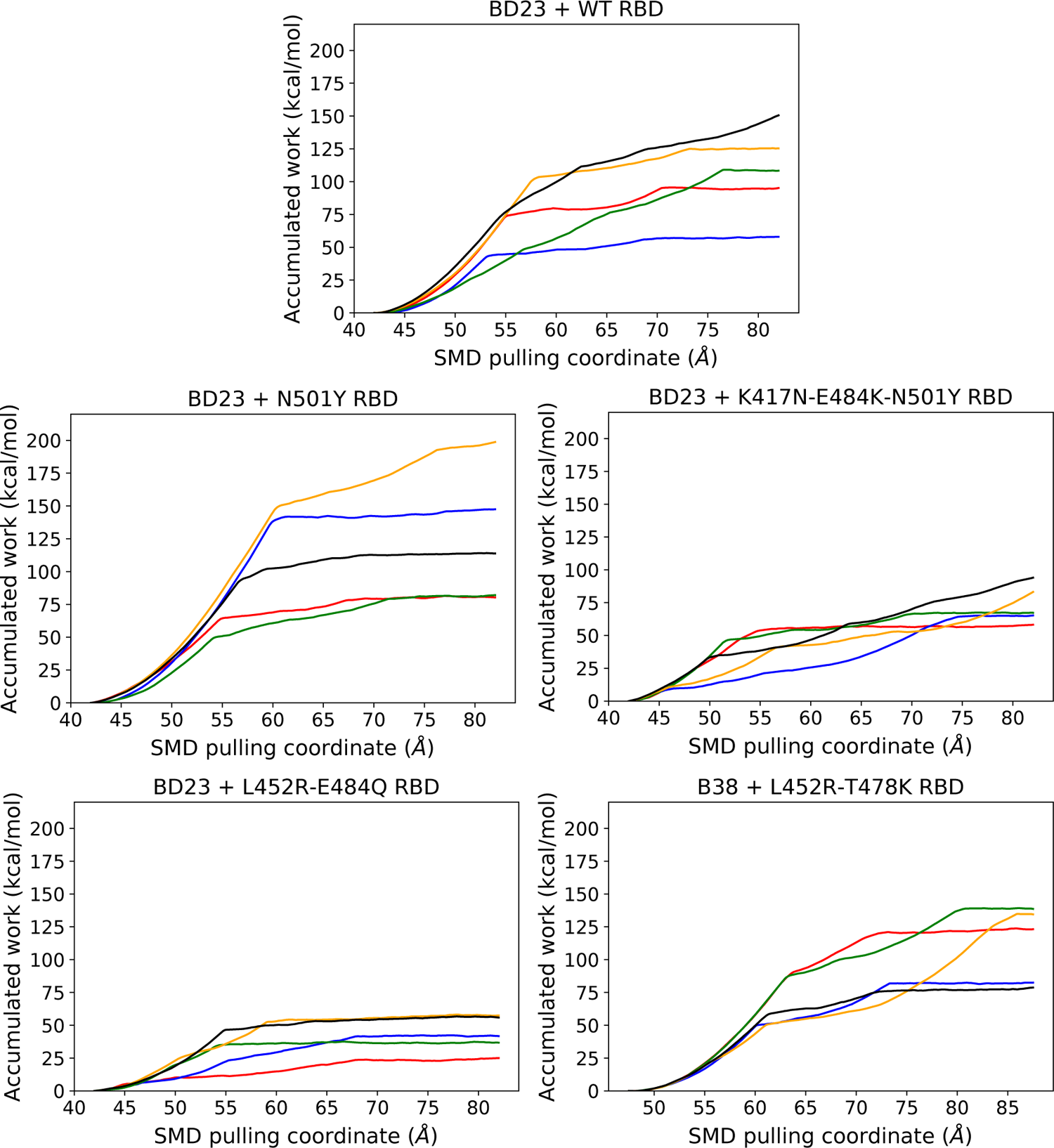
Identical to Figure S18, except for the antibody BD23 Fab.

**Figure S20:**
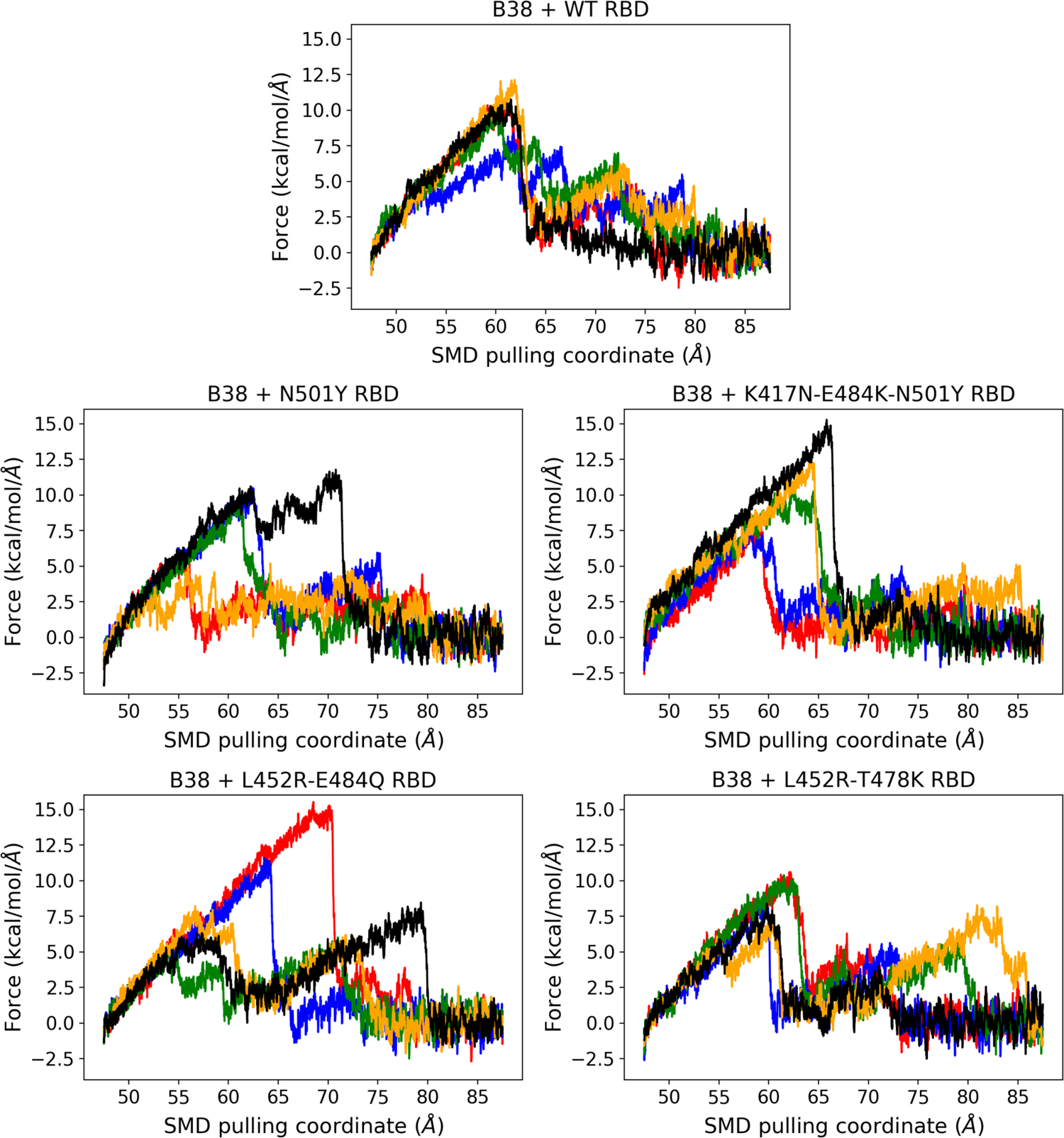
The pulling force required to move the B38 antibody away from the WT and the mutant RBDs along the center of mass distance between RBD and Ab. The color scheme is same as Fig. S18

**Figure S21:**
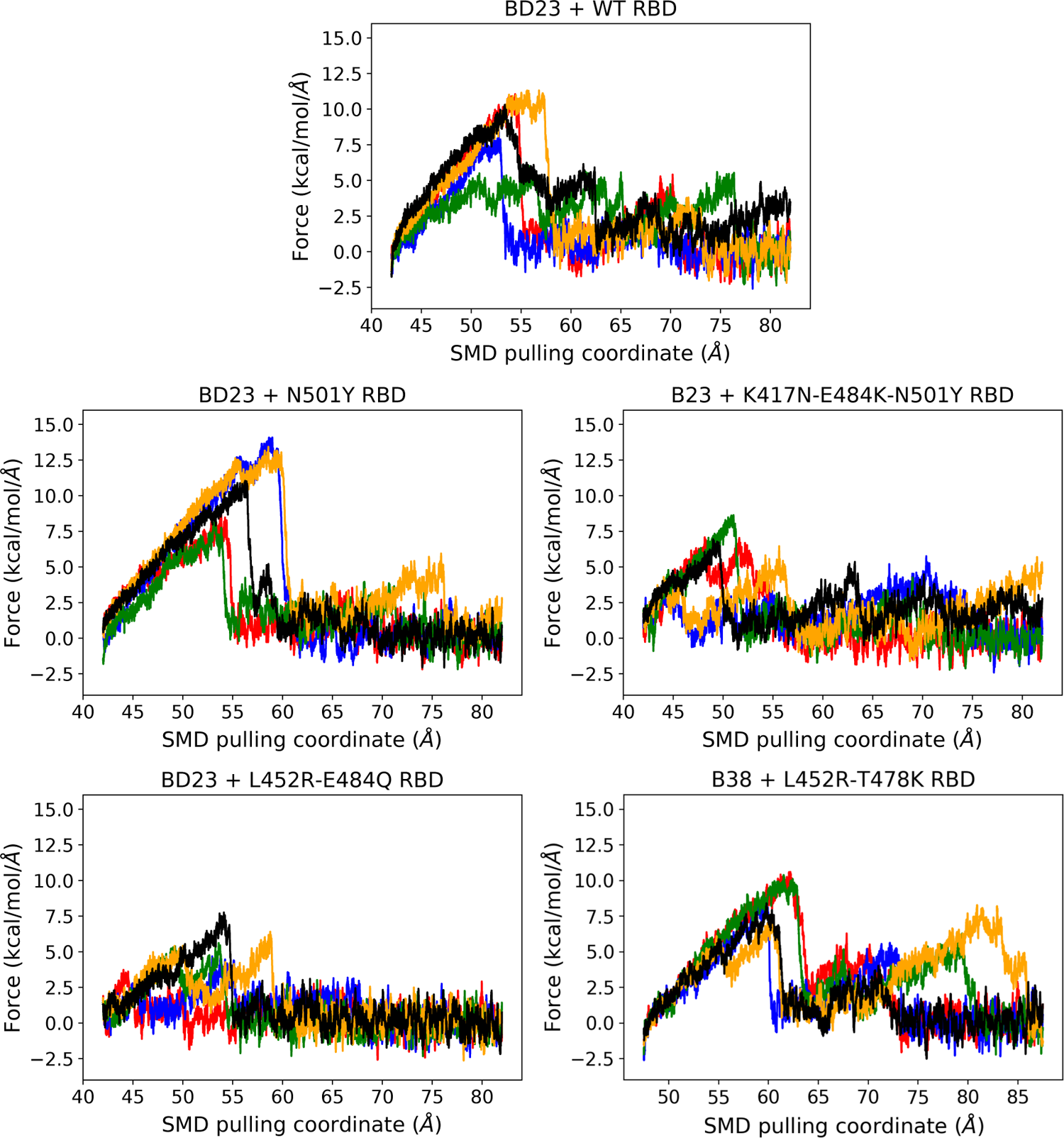
Identical to Figure S20, except for the antibody BD23 Fab.

**Figure S22:**
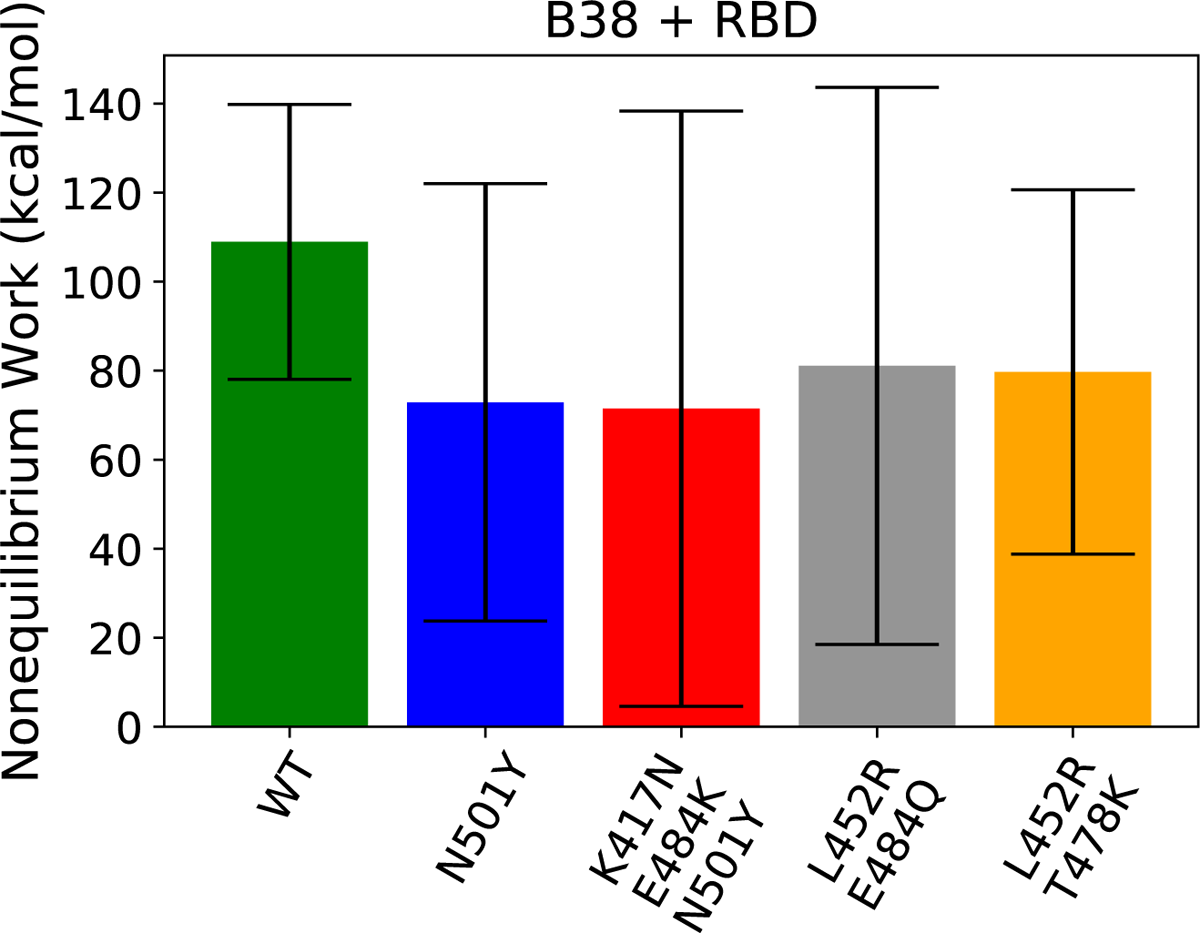
Average non-equilibrium work for the dissociation of the B38 antibody from the RBD, along with error bars (root mean squared errors).

**Figure S23:**
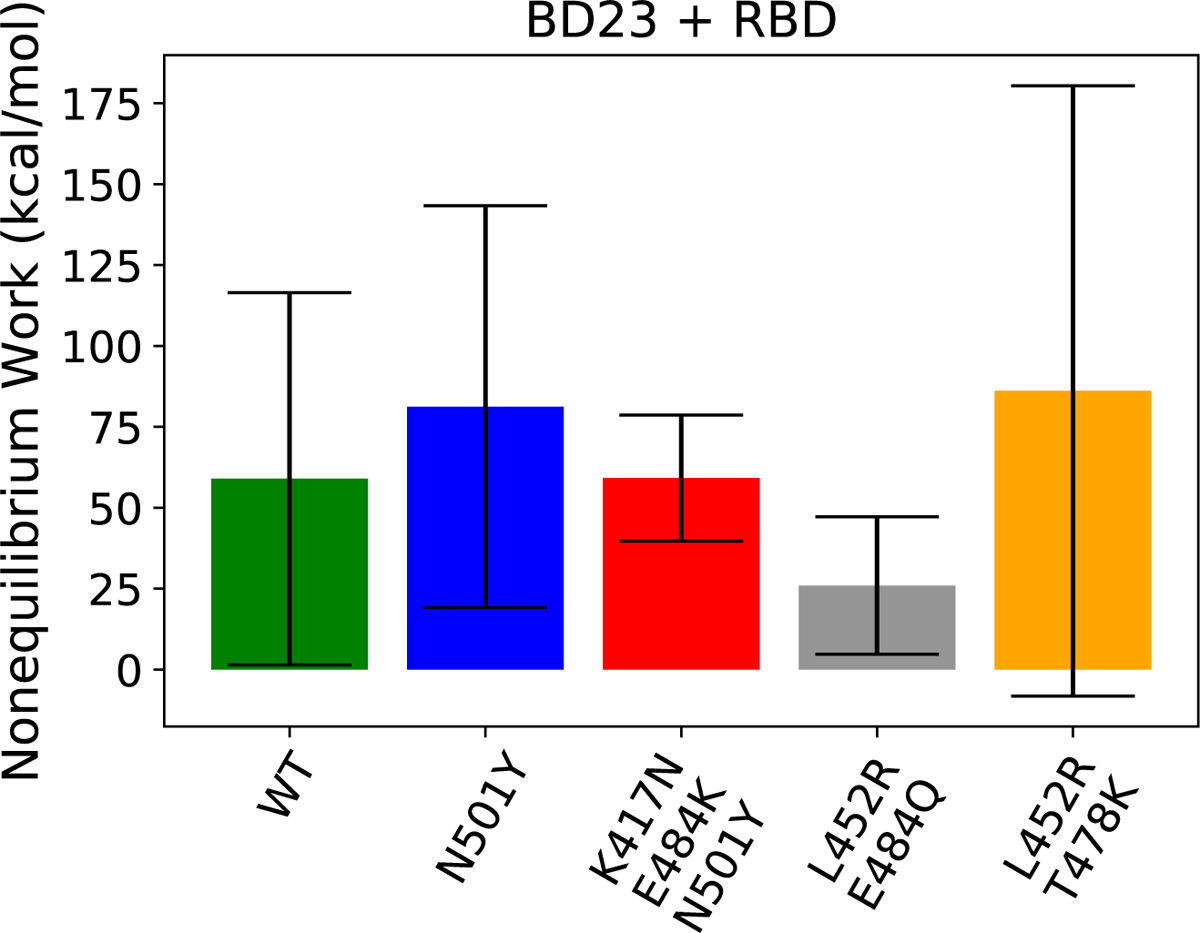
Average non-equilibrium work for the dissociation of the BD23 antibody from the RBD, along with error bars (root mean squared errors).

**Figure S24:**
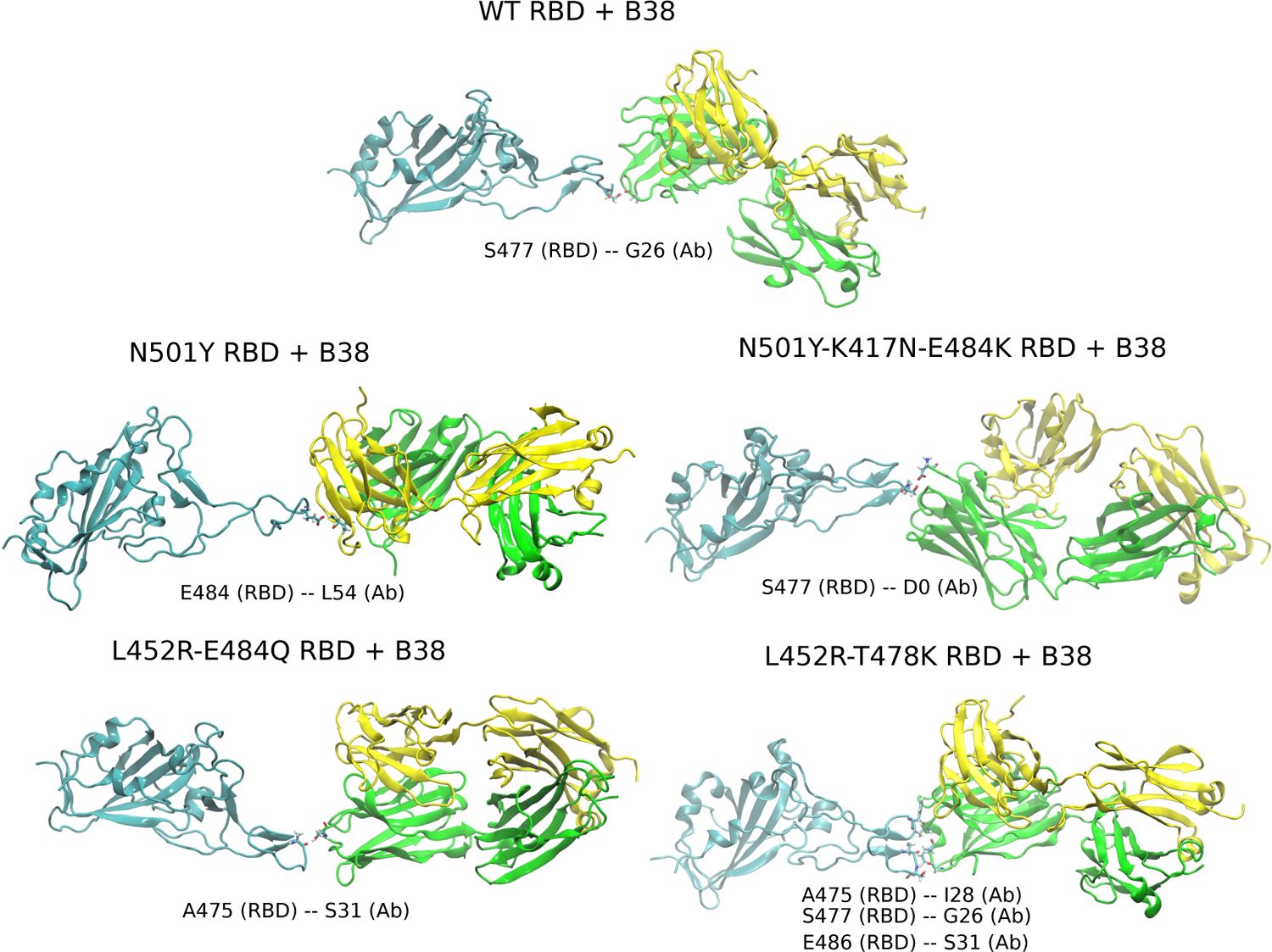
The intermediate structures of the B38 dissociation from the RBD of different variants, sampled from the lowest work SMD trajectory for each respective system. The figure depicts the last intact hydrogen bond between RBD and antibody during the dissociation process.

**Figure S25:**
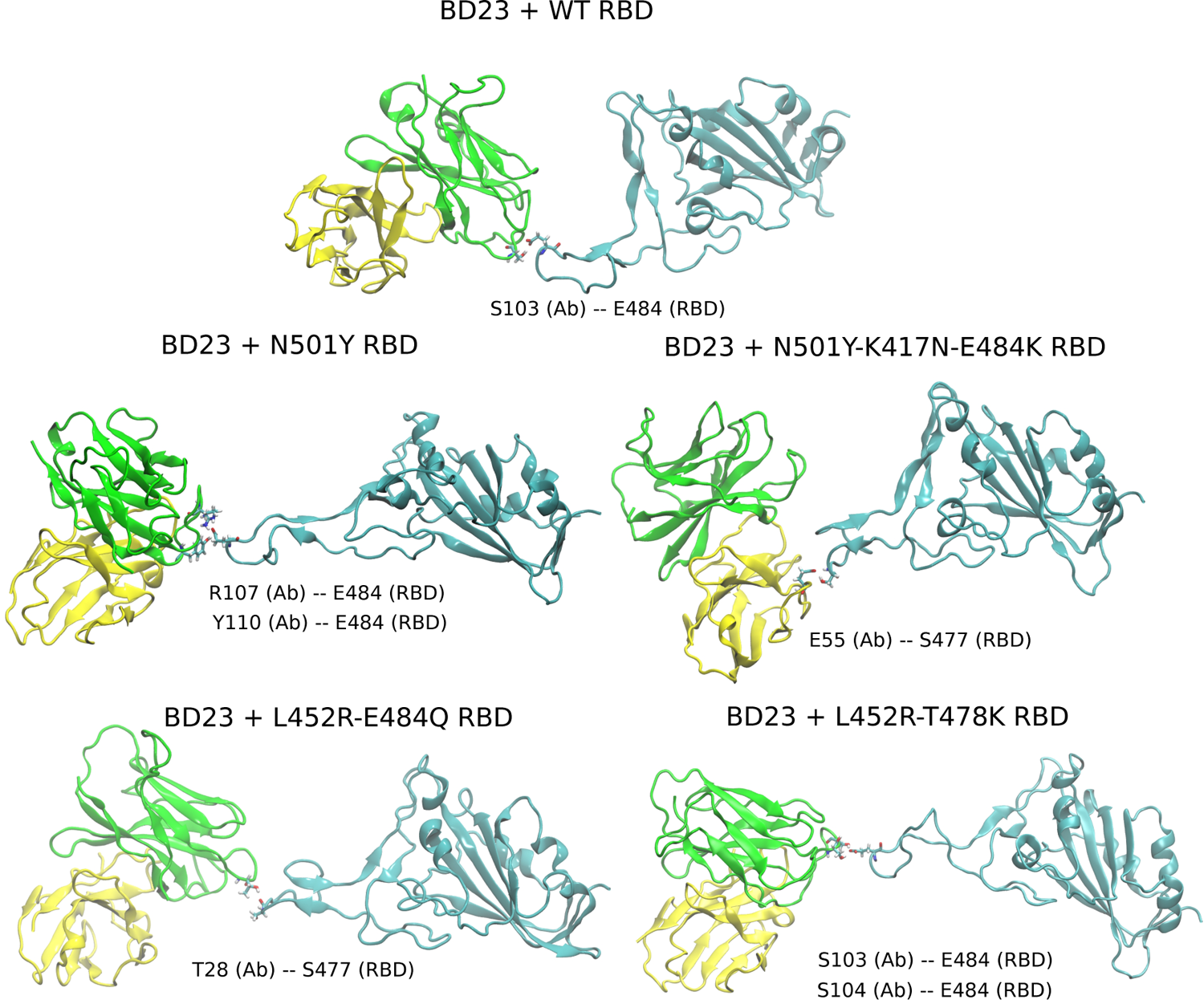
Identical to Figure S24, except for the antibody BD23 Fab.

**Figure S26:**
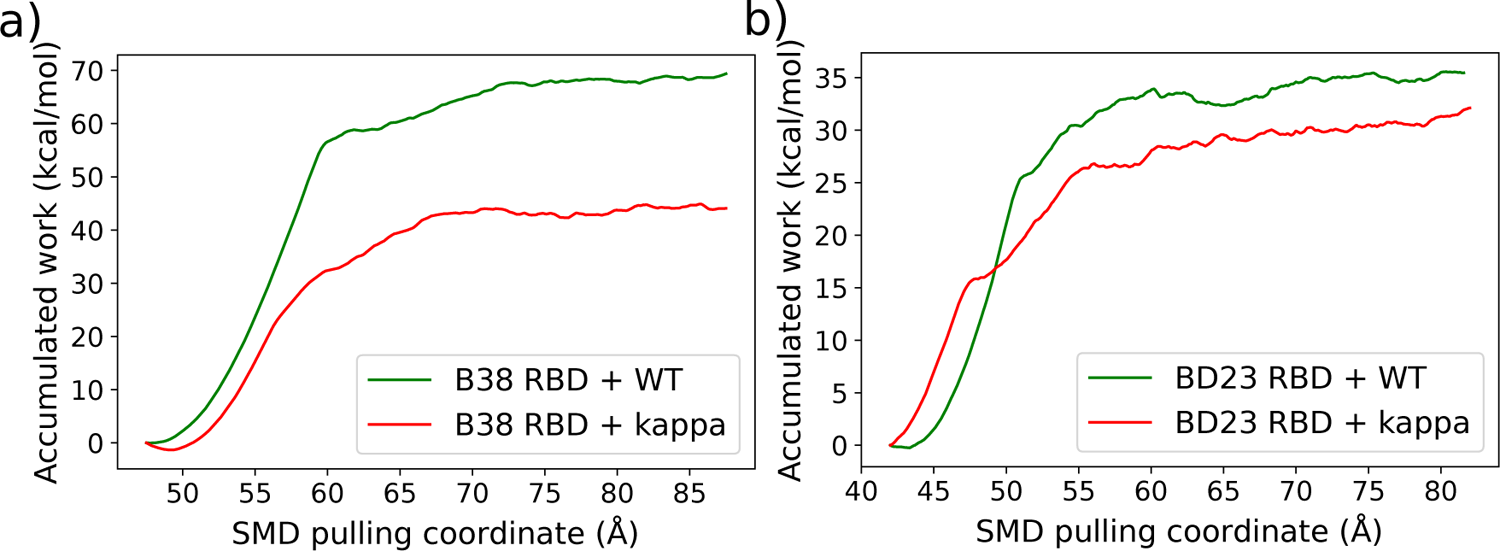
The work required to pull the (a) B38 antibody or the (b) BD23 antibody, along the center of mass distance between the RBD and antibody using *explicit solvent* SMD simulation.

**Figure S27:**
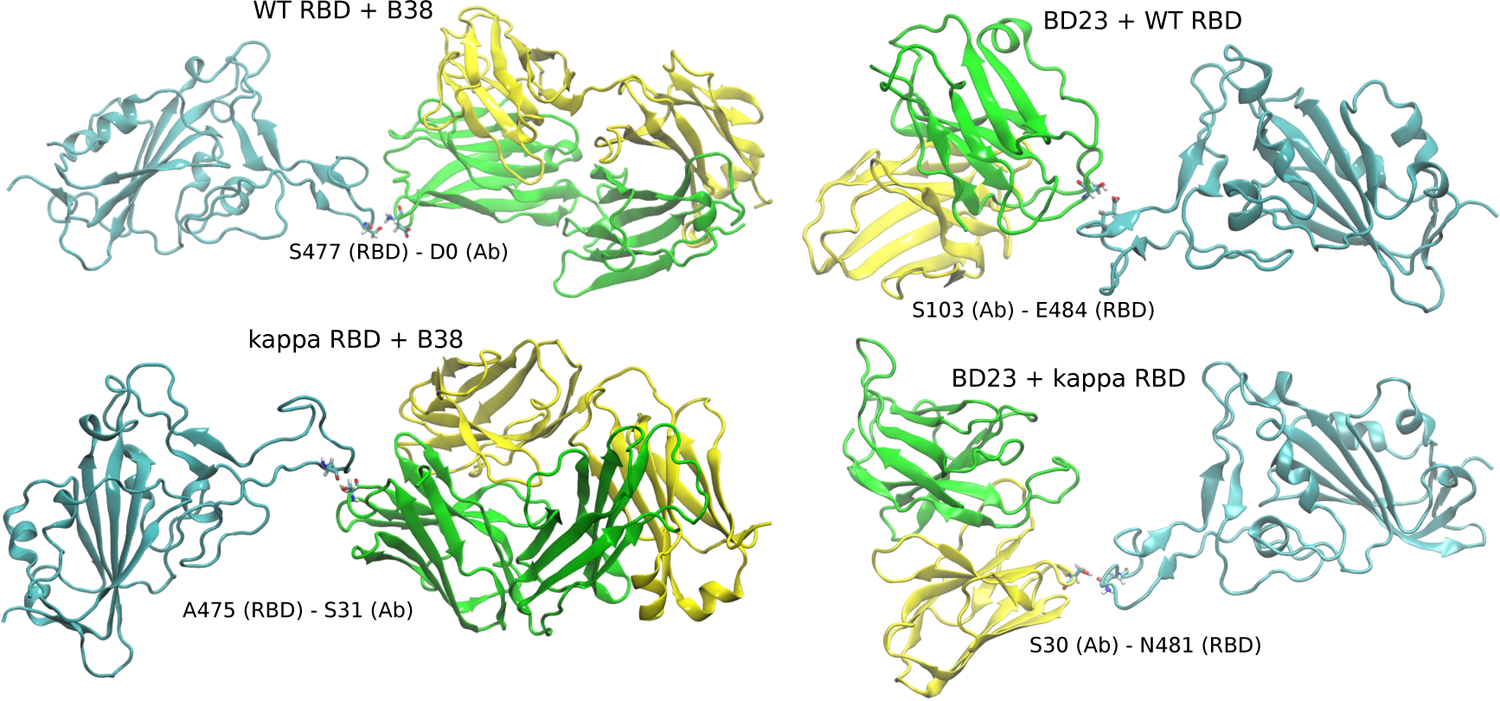
The intermediate structures for the dissociation of the B38 and the BD23 antibodies from the RBD of the WT and the *kappa* variants of the SARS-CoV-2 spike protein, in explicit solvent. The water and ions are not shown for clarity.

## Notes

https://doi.org/10.5281/zenodo.5676393

https://github.com/dhimanray/COVID-variant-antibody.git

### Competing Interest Statement

The authors have declared no competing interest.

